# Sparse genetic tracing reveals regionally specific functional organization of mammalian nociceptors

**DOI:** 10.1101/149641

**Authors:** William Olson, Ishmail Abdus-Saboor, Lian Cui, Justin Burdge, Tobias Raabe, Minghong Ma, Wenqin Luo

## Abstract

The human distal limbs have a high spatial acuity for noxious stimuli but a low density of pain-sensing neurites. To elucidate mechanisms underlying the ‘pain fovea’, we sparsely traced non-peptidergic nociceptors across the body using a newly generated *MrgprD^CreERT2^* mouse line. We found that mouse plantar paw skin also has a low density of MrgprD^+^ neurites, and individual arbors in different locations are comparable in size. Surprisingly, the central arbors of plantar paw and trunk innervating nociceptors have distinct morphologies in the spinal cord. This regional difference is well correlated with a heightened signal transmission for plantar paw circuits, as revealed by both spinal cord slice recordings and behavior assays. Taken together, our results elucidate a novel somatotopic functional organization of the mammalian pain system and suggest that regional central arbor structure could facilitate the magnification of plantar paw regions to contribute to the ‘pain fovea’.

## Introduction

The skin mediates physical contact with environmental mechanical, thermal, and chemical stimuli. As an animal moves through the world, certain parts of the skin are likely sites of “first contact” with these stimuli (for instance, the distal limbs, face/whiskers, and tail for quadrupedal mammals like mice). Therefore, these somatosensory regions require heightened sensitivity to fulfil behaviorally relevant functions, such as environment exploration.

Touch and pain are the two most important types of somatosensation for this functional purpose: touch allows for feature detection while pain prevents tissue damage. Indeed, classic work has defined important regional specialization of the nervous system for tactile sensation in these areas. Two mechanisms in the peripheral organization of the discriminative touch system facilitate high spatial acuity sensation in the primate distal limbs and mouse whisker pad. These are the *high innervation density* and *smaller receptive field sizes* of the primary light touch neurons, the Aβ mechanoreceptors, in these regions^1-7^. In contrast, the question of whether regional specialization exists in the mammalian pain system has remained elusive until recently. Upon stimulation using nociceptive-specific laser beams, human subjects show a heightened spatial acuity in the distal limbs (especially the fingertips) for pain stimuli, much like they do for touch stimuli^8-10^. This suggests that this region is also a “pain fovea”. However, human fingertip skin has a relatively *low density of pain-sensing neurites*^9^. While this suggests that region-specific organization likely exists in pain circuits downstream of the periphery (i.e. central nervous system), currently the exact neural mechanisms of the pain fovea are unclear.

Pain stimuli are detected by primary sensory neurons called nociceptors, which have cell bodies in the dorsal root ganglia (DRG) or trigeminal ganglia (TG) and axons that bifurcate into peripheral and central branches. The peripheral projection of a nociceptor usually terminates as a free nerve arbor in the skin or deeper tissues, while the central projection terminates in an arbor in the dorsal horn (DH) of the spinal cord or caudal medulla. Though previous work has mapped the peripheral receptive fields or traced the central terminals of mammalian nociceptors^11-21^, these studies have not established a model for the somatotopic functional organization of the mammalian pain system due to the limited number of neurons traced from restricted skin regions.

We therefore sought to reveal the region-specific organization of mammalian nociceptors across the entire somatotopic map. We generated a novel *MrgprD^CreERT2^* mouse line to perform systematic sparse genetic tracing of a population of non-peptidergic nociceptors that mediate mechanical pain and beta-alanine (B-AL) triggered itch^22, 23^. We chose these nociceptors because they are the most abundant type of cutaneous nociceptor and they likely correspond to the main type of free nerve terminals stained with anti-PGP9.5 antibody in previous human skin biopsy data^9, 24^.

Indeed, like the human skin biopsy results, we found that MrgprD^+^ neurites have a comparatively low neurite density in plantar paw compared to trunk skin. In addition, and in contrast to the Aß mechanoreceptors, sparse genetic tracing revealed that the arbor field sizes of individual nociceptors are comparable between different skin regions, whereas plantar paw and trunk innervating nociceptors display distinct morphologies in their central terminals. Lastly, using *MrgprD^CreERT2^*; *Rosa^ChR2-EYFP^* mice, we specifically activated these nociceptors using blue laser light during *in vitro* spinal cord slice recordings and during behavior assays. We found that, while almost all layer II DH neurons in all locations receive direct MrgprD^+^ afferent input, the optical threshold required to induce postsynaptic responses is much lower in plantar paw regions. This was paralleled by a decrease in the light intensity threshold required to elicit a withdrawal response in paw, compared to upper thigh, skin stimulation. Collectively, we have identified a previously unappreciated somatotopic difference in the functional organization of mammalian nociceptors. Our anatomical, physiological, and behavior data suggest that region-specific central arbor structure could help to magnify the representation of plantar paw nociceptors in the DH to establish the ‘pain fovea’.

## Results

### Generation and specificity of *MrgprD^CreERT2^* mice

Given that previous descriptions of nociceptor structure have not allowed for systematic comparisons between body regions, we sought to use sparse genetic labeling to trace single nociceptor morphologies across the entire somatotopic map. We generated a mouse line in which a tamoxifen-inducible Cre (CreERT2) cassette is knocked into the coding region of Mas-related gene product receptor D (*MrgprD*) (Figure 1A, Figure S1). Consistent with the previous finding that *MrgprD* is expressed more broadly in early development than in adulthood^25^, early embryonic (E16.5-E17.5) tamoxifen treatment of *MrgprD^CreERT2^* mice labels MrgprD^+^ neurons along with non-peptidergic neurons expressing other *Mrgpr* genes, such as *MrgprA3* and *MrgprB4* (Figure S2). In contrast, when we crossed *MrgprD^CreERT2^* mice with a Cre-dependent *Rosa^ChR2-EYFP^* line and provided postnatal (P10-P17) tamoxifen treatment (Figure 1B), MrgprD^+^ non-peptidergic nociceptors were specifically labeled. We examined these treated mice at 4 postnatal weeks or older (>4pw), a time point at which MrgprD^+^ non-peptidergic nociceptors have completely segregated from other Mrgpr^+^ DRG neurons^25^. We found that ChR2-EYFP^+^ DRG neurons bind IB4 (a marker for non-peptidergic DRG neurons) but do not express CGRP (a marker for peptidergic DRG neurons)^24^ (Figure 1C), and ChR2-EYFP^+^ DH terminals similarly overlap with IB4 but not CGRP (Figure 1G). Double *in situ* hybridization demonstrated that this strategy efficiently labels DRG neurons expressing *MrgprD* (88.1 ± 1% of ChR2-EYFP^+^ neurons, *n* = 3 animals) but not those expressing *MrgprA3* (1.4 ± 0.1%) or *MrgprB4* (0.4 ± 0.3%) (Figure 1D-F). Almost all *MrgprD* expressing neurons were labeled with ChR2-EYFP (92.9 ± 4.6% of *MrgprD*^+^ neurons) by this treatment. Therefore, this newly generated inducible *MrgprD^CreERT2^* line allows for the specific and efficient targeting of adult MrgprD^+^ nociceptors.

We then sought to trace individual MrgprD+ non-peptidergic nociceptors using sparse genetic labeling. When crossed with a Cre-dependent alkaline phosphatase reporter line (*Rosa^iAP^*), we found that sparse recombination occurs in the absence of tamoxifen treatment (Figure 2A&B). This background recombination labels 3-11 neurons/DRG (5.2 ± 1.6 neurons/DRG, *n* = 47 DRGs from 3 animals) in 3-4 pw animals (Figure 2C), which represents <1% of the total MrgprD^+^ nociceptor population^24, 26, 27^. The sparsely labeled DRG neurons co-express non-peptidergic nociceptor markers peripherin, PAP^28^, and RET (Figure 2D, F, H), but do not express NF200 or CGRP (Figure 2E, G). To further determine the specificity of this sparse recombination, we used an *MrgprD^EGFPf^* knock-in line^24^, in which expression of EGFP mimics the dynamic expression of endogenous *MrgprD*. We generated *MrgprD^CreERT2/EGFPf^*; *Rosa^iAP^* mice and found that almost all AP^+^ neurons co-express *MrgprD^EGFPf^* (93. 7 ± 2.3%, *n* = 126 AP^+^ neurons from 3 animals) (Figure 2I) in 3 to 4pw mice. This result indicates that, although *MrgprD* is broadly expressed during early development, this background recombination occurred preferentially in adult MrgprD^+^ nociceptors.

**Figure 1.**
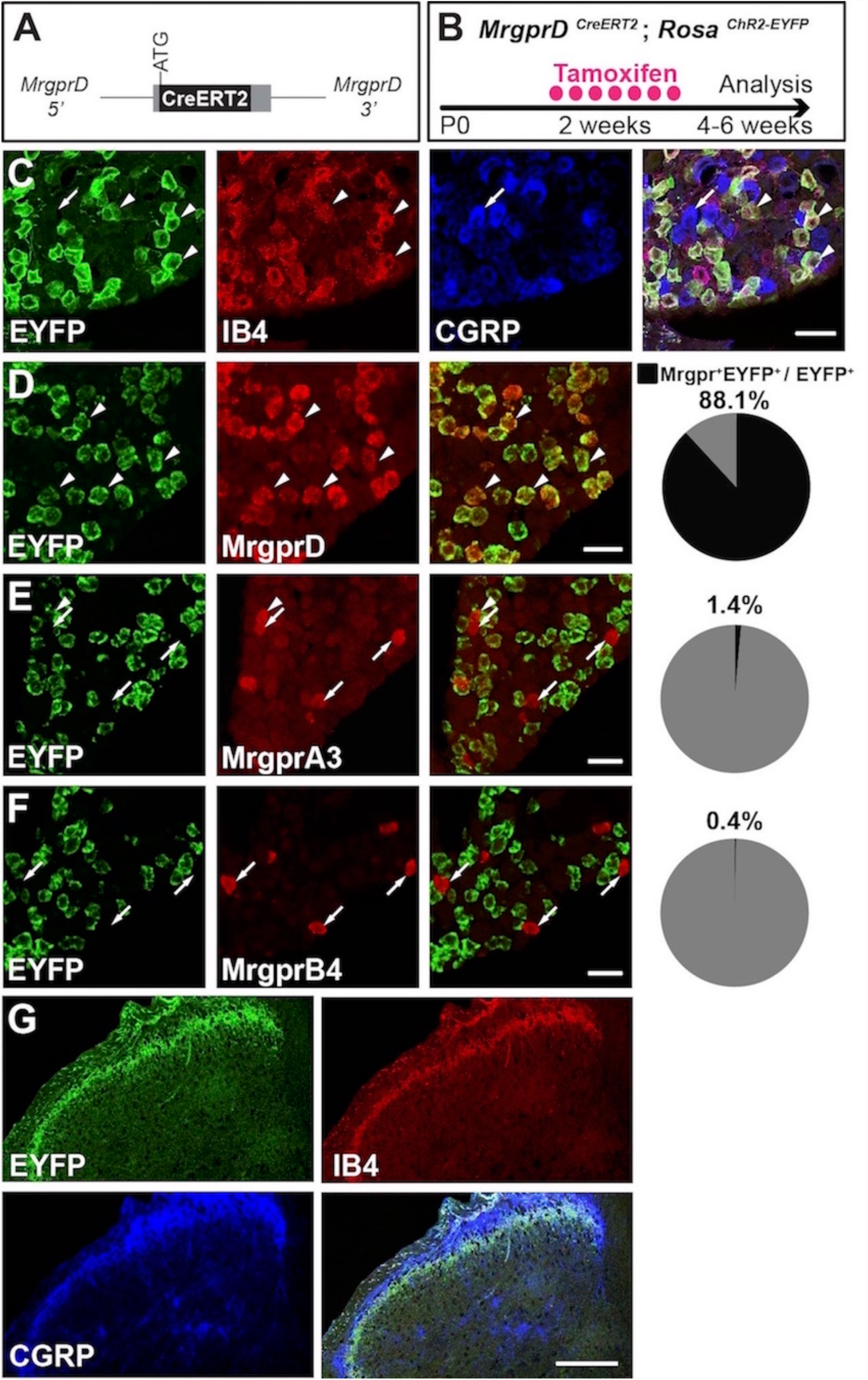
*MrgprD^CreERT2^* mice can mediate recombination specifically in adult MrgprD^+^ non-peptidergic nociceptors. (A) Knock-in *MrgprD^CreERT2^* allele. (B) Illustration showing tamoxifen treatment scheme, 0.5 mg tamoxifen / day, P10-P17 treatment of *MrgprD^CreERT2^*; *Rosa^ChR2-EYFP^* mice. (C) Triple staining of DRG section showing EYFP overlaps with IB4 but not CGRP. (D-F) Double fluorescent *in situ* DRG sections showing EYFP in *MrgprD* (D) but not *MrgprA3* (E) or *MrgprB4* (F) cells. Pie charts show overlap quantification (% of EYFP^+^ cells that co-express Mrgpr, *n* = 3 animals). (G) DH section showing EYFP^+^ terminal overlap with IB4 but not CGRP. Arrowheads show overlapping cells, arrows show non-overlapping cells. Scale bars = 50μm (C-F), 100μm (G).

**Figure 2.**
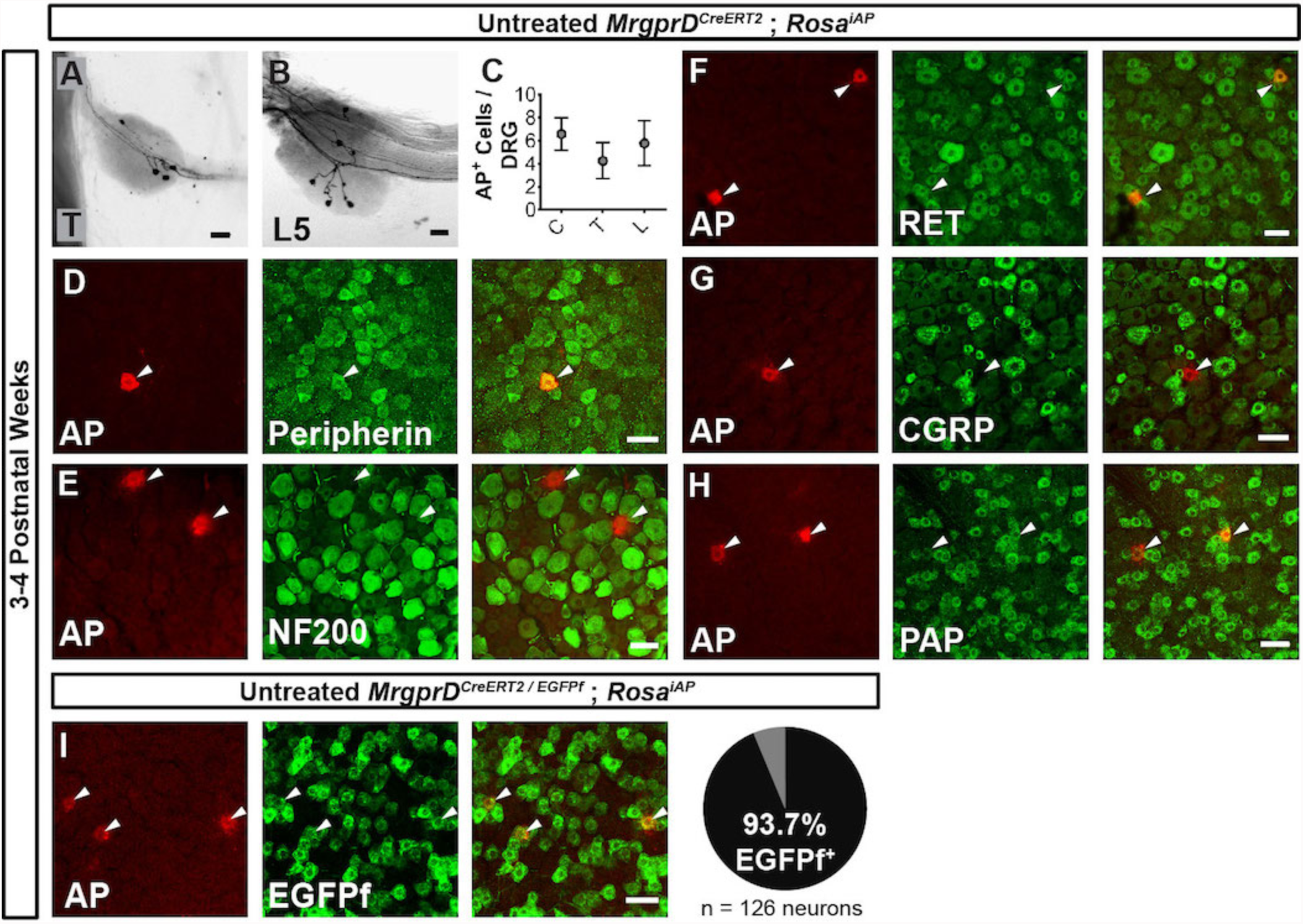
Sparse MrgprD^+^ nociceptor labeling in untreated 3-4 pw *MrgprD^CreERT2^*; *Rosa^iAP^* mice. (A©B) Whole mount AP DRG staining of thoracic (A) and L5 (B) DRGs. (C) AP^+^ cells / DRG for cervical (C), thoracic (T), and lumbar (L) DRGs, *n* = 47 DRGs from 3 animals. (D-H) Whole mount DRG immunostaining plus AP fluorescent staining. Sparse AP^+^ cells express non-peptidergic nociceptor markers peripherin (D), RET (F) and PAP (H) but not large diameter neuron maker NF200 (E) or peptidergic marker CGRP (G). (I) Whole mount EGFPf immunostaining plus AP fluorescence staining of untreated *MrgprD^CreERT2/EGFPf^*; *Rosa^iAP^* DRGs. AP^+^ neurons are MrgprD^+^ nociceptors. Quantification of overlap (% of AP^+^ cells that co-express *MrgprD^EGFPf^*, *n* = 126 neurons from 3 animals). Scale bars = 50μm.

### Genetic tracing of MrgprD^+^ skin terminals reveals a relatively comparable organization in the periphery

MrgprD^+^ neurons innervate both hairy and glabrous skin and are the most abundant class of cutaneous free nerve arbors^24^. To systematically compare the peripheral single-cell structure of mammalian pain neurons across the somatotopic map, we performed whole mount colorimetric AP staining using untreated 3-4 pw *MrgprD^CreERT2^*; *Rosa^iAP^* skin.

We found that 98.4% (130/132 arbors, *n* = 4 animals) of single-cell arbors have a “bushy ending” morphology (Figure 3A&B, D&E, Figure S3A)^29^, featuring thickened terminal structures in the epidermis. The distal ends of arbors in glabrous plantar paw skin have single, un-branched thickened neurites (Figure 3A&B), while arbors in the hairy skin feature both un-branched neurites as well as dense neurite clusters (Figure 3D&E). These dense clusters are seen in all analyzed regions that have hairy skin and likely innervate the necks of hair follicles (red arrowheads in Figure 3E&F) (see also Figure S5C)^24^. A very small minority (1.6%) of arbors in the hairy skin have “free endings”^29^ lacking these thickened structures (Figure S3A-C).

**Figure 3.**
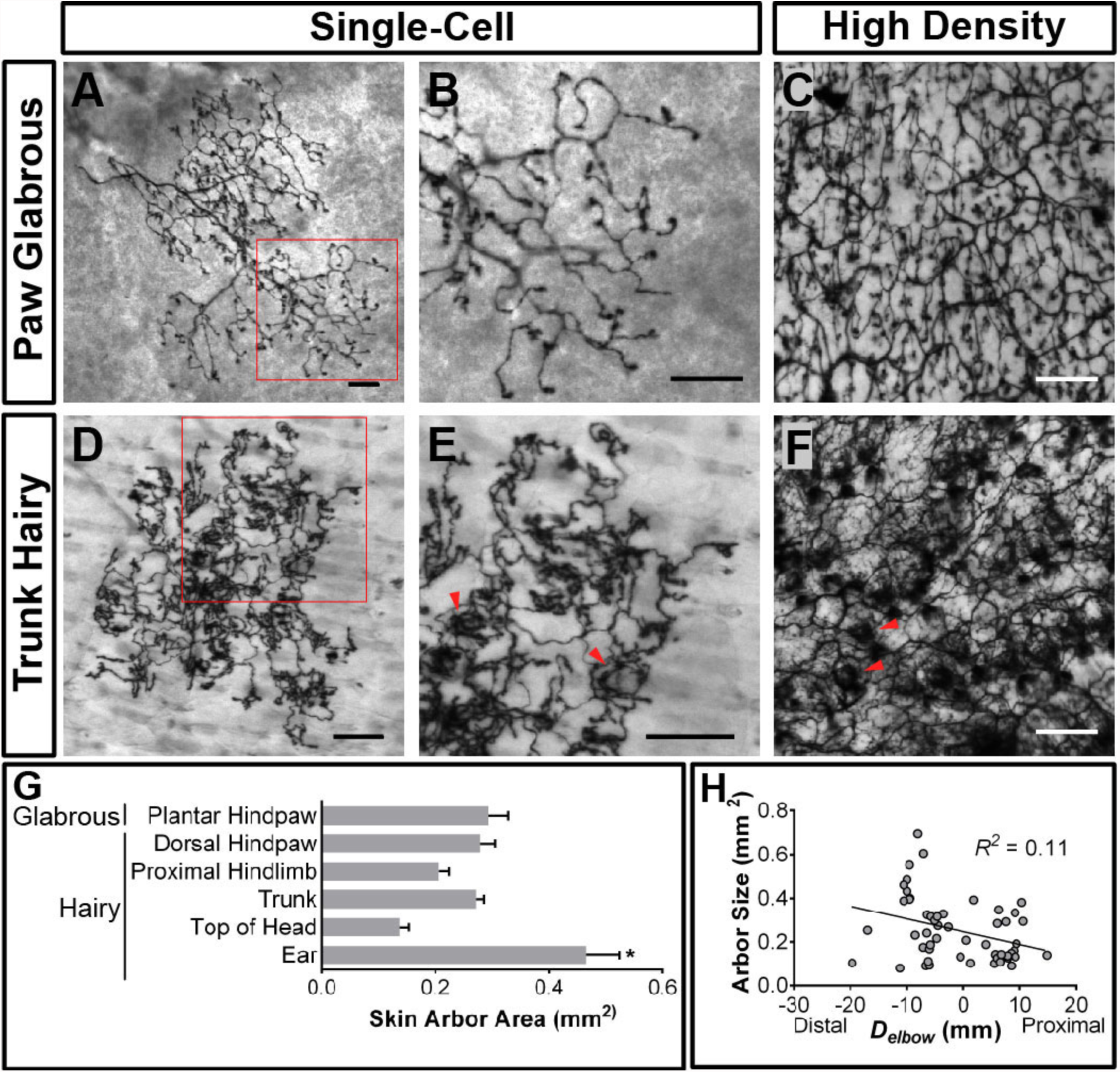
Peripheral organization of non-peptidergic nociceptors in 3-4 pw *MrgprD^CreERT2^*; *Rosa^iAP^* mice. (A, B, D, E) Sparse labeled non-peptidergic nociceptors show bushy-ending structure in the glabrous skin (A-B) and trunk hairy skin (D-E). B, E, high magnification images of regions boxed in A and D, respectively. (C, F) High-density labeled (0.5mg tamoxifen at P11) glabrous and hairy skin. Overall neurite densities are lower in glabrous compared to hairy skin. Red arrowheads in E&F mark neurite clumps that likely surround hair follicles. (G) Arbor areas in different skin regions. *n* = 173 terminals from 9 animals. * = *p*<0.05 (one-way ANOVA with Tukey’s multiple comparisons test). (H) Arbor areas in the hind limb skin vs. proximodistal distance (*D_elbow_*) from the elbow (point 0, terminals distal to this edge are given negative *D_elbow_* values). *n* = 52 arbors from 4 animals. No clear relationship between proximodistal location and size is evident. Scale bars = 100μm.

MrgprD^+^ non-peptidergic nociceptive field sizes range from 0.08 to 0.9 mm^2^, with the smallest average field size found in the head skin between the ears and in the proximal limbs (Table 1, Figure 3G). Interestingly, non-peptidergic nociceptors innervating the distal limbs (plantar and dorsal paw skin) have average arbor sizes close the middle of this range, and distal limb and trunk arbors are comparable in size (Table 1, Figure 3G&H). In addition, consistent with human skin^8^, whole-population labeling of MrgprD^+^ fibers using tamoxifen (0.5 mg at P11) reveals that the overall neurite density is slightly lower in the paw glabrous skin compared to trunk hairy skin (Figure 3C&F). In short, in contrast to mammalian Aβ mechanoreceptors, plantar paw innervating mouse non-peptidergic nociceptors do not display higher neurite density or smaller receptive field sizes.

**Table 1.** Summary of peripheral terminals of sparsely labeled MrgprD^+^ non-peptidergic nociceptors. Data pooled from seven 3pw animals. Asterisk (*) indicates a significant difference (*p*<0.05, One-way ANOVA with Tukey’s multiple comparison test) in average size in pair-wise comparisons with all other regions.

| **Region** | **n** | **Avg area**<br>**(mm<sup>2</sup>)** | **SE** | **Min area**<br>**(mm<sup>2</sup>)** | **Max area**<br>**(mm<sup>2</sup>)** |
| --- | --- | --- | --- | --- | --- |
| Plantar hindpaw<br>(glabrous) | 22 | **0.29** | 0.03 | 0.08 | 0.86 |
| Dorsal hindpaw | 27 | **0.28** | 0.03 | 0.15 | 0.54 |
| Proximal hindlimb | 29 | **0.21** | 0.02 | 0.14 | 0.32 |
| Dorsal forepaw | 9 | **0.26** | 0.07 | 0.11 | 0.45 |
| Proximal forelimb | 3 | **0.14** | 0.01 | 0.13 | 0.14 |
| Trunk | 77 | **0.27** | 0.01 | 0.10 | 0.9 |
| Top of head | 6 | **0.14** | 0.02 | 0.10 | 0.20 |
| Ear* | 11 | **0.47** | 0.06 | 0.22 | 0.93 |

### MrgprD^+^ nociceptors show regionally distinct organization in their central arbors

Since the peripheral organization of MrgprD^+^ non-peptidergic nociceptors does not exhibit an obvious mechanism to facilitate heightened sensitivity in the plantar paw, we next used whole mount AP staining of untreated 3-4 pw *MrgprD^CreERT2^*; *Rosa^iAP^* spinal cords to compare their central arbors between regions. Non-peptidergic nociceptor central branches enter the spinal cord through the dorsal root, travel rostrally or caudally for 0 to 3 segments, and then dive ventrally to establish arbors in layer II of the DH (Table 2)^24^. Most MrgprD^+^ central branches do not bifurcate (65.8%, *n* = 234 neurons from 3 animals), and most also terminate within the segment of entry (72.6%) (Table 2). However, some (34.2%) bifurcate one or more times in the spinal cord, and some (27.3%) travel up to 3 segments from the point of entry (Table 2). For the central branches that bifurcate, most of their secondary/tertiary branches join other branches from the same neuron to co-form one axonal arbor, while some end with a second arbor or terminate in the spinal cord without growing an arbor (Table 2, Figure S4A&B). The majority (91.9%) of labeled nociceptors have only one arbor, but a few have no (0.4%), 2 (6.8%), or 3 (0.9%) central arbors (Table 2).

**Table 2.** Summary of central innervation patterns of sparsely labeled MrgprD^+^ non-peptidergic nociceptors. Data pooled from three 3pw animals.

|  | *n* | % of Total |
| --- | --- | --- |
| Total neurons | 234 |  |
| Segments traveled from point of entry |  |  |
| 0 | 170 | 72.6 |
| 1 | 54 | 23.1 |
| 2 | 9 | 3.8 |
| 3 | 1 | 0.4 |
| Direction traveled (for axons traveling 1-3 segments) |  |  |
| Caudal | 46 | 19.7 |
| Rostral | 18 | 7.7 |
| No central branch bifurcations | 154 | 65.8 |
| 1 central branch bifurcation | 72 | 30.8 |
| >1 central branch bifurcation | 8 | 3.4 |
| No central terminals | 1 | 0.4 |
| 1 central terminals | 215 | 91.9 |
| 2 central terminals | 16 | 6.8 |
| 3 central terminals | 2 | 0.9 |

Strikingly, MrgprD^+^ non-peptidergic nociceptor central arbors display two different morphologies that can be distinguished by the ratio of their mediolateral width to their rostrocaudal height (W/H ratio) (Figure 4H). We defined arbors with W/H ratios >0.2 as “round” (Figure 4A-C, H, and Table 3) and arbors with W/H ratios <0.2 as “long and thin” (Figure 4B-F, H, and Table 3). Further, these morphological types are regionally segregated. Round arbors are found in DH regions known to represent the distal limbs (medial cervical and lumbar enlargements) as well as tail, anogenital (sacral DH), and head/face (descending trigeminal terminals in the upper cervical DH and medulla) skin (Figure 4B&C, H, J, and Figure S4C)^30, 31^. Long arbors are instead located in regions corresponding to the proximal limbs (lateral cervical and lumbar enlargements) and trunk skin (thoracic DH) (Figure 4B-F, H, J)^30-32^. While round and long arbors differ in morphology, they do not differ in area (Figure 4G).

**Figure 4.**
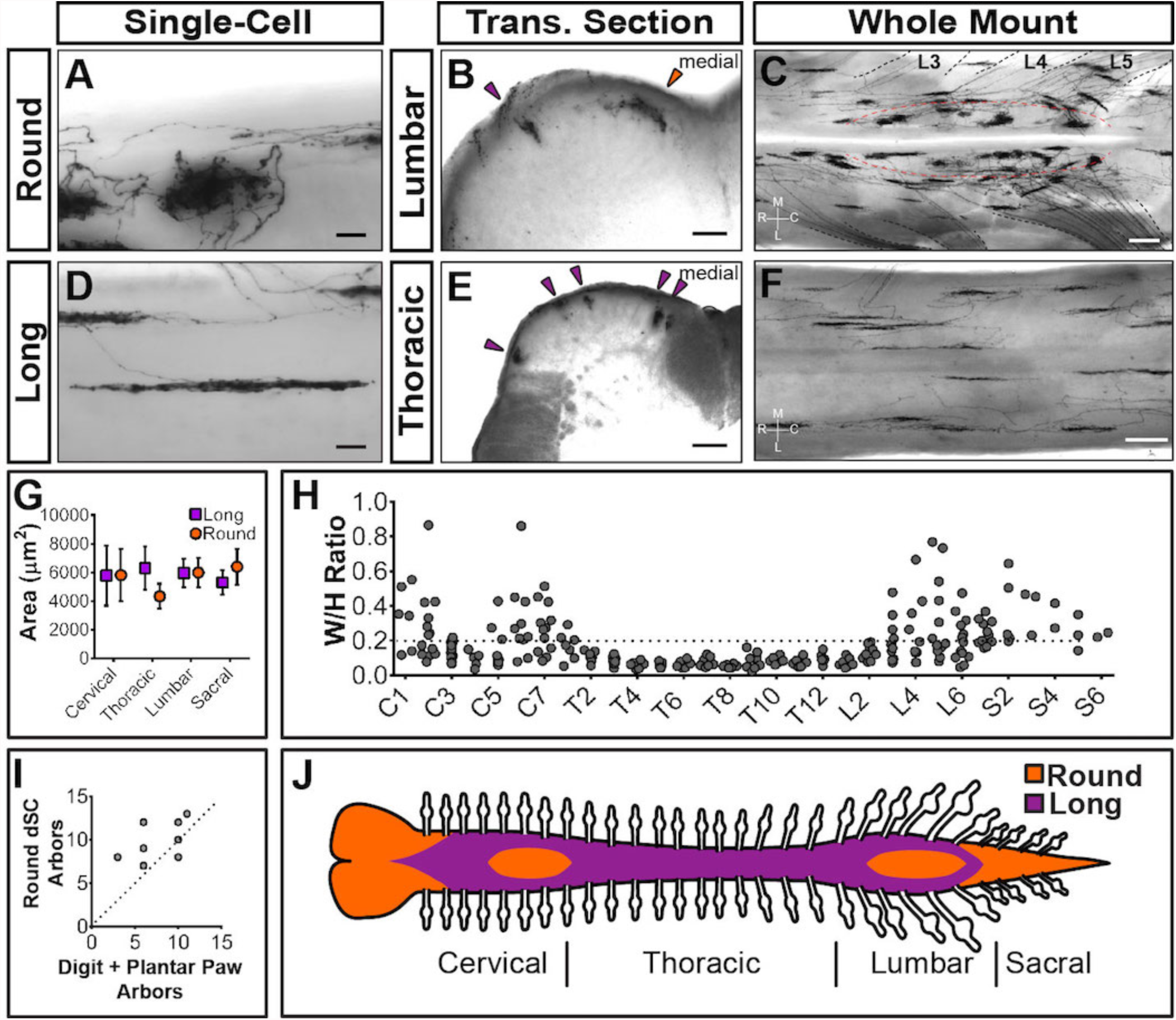
Sparsely labeled *MrgprD^CreERT2^*; *Rosa^iAP^* nociceptors have region-specific central arbor morphologies. (A-F) Round and long non-peptidergic central arbors seen in top-down whole mount (A, C, D, F) and transverse section (B&E) spinal cords. Round arbors are in the medial lumbar enlargement (B&C) while long arbors are in the lateral lumbar enlargement and thoracic spinal cord (B-F). Dorsal roots are outlined and labeled in C. Red dashed lines outline the round arbor zone. Orange arrowhead marks a round terminal, purple arrowheads mark long terminals. M, medial. L, lateral. R, rostral. C, caudal. (G) Round and long (defined by ratio in H) arbor areas are comparable for all regions. (H) Arbor Width/Height ratios by ganglion of origin. Round terminals: W/H > 0.2. *n =*368 terminals from 7 animals (I) Comparison of the number of labeled arbors in the hindlimb digit and plantar paw skin with the number of ipsilateral round terminals in the dorsal horn. *n* = 4 animals, dotted line shows 1:1 relationship. (J) Illustration showing the distribution of round (orange zone) and long (purple zone) arbors in the spinal cord. Scale bars = 50μm (A&B, D&E), 250μm (C&F).

**Table 3.** Summary of non-peptidergic nociceptor central terminal height and width measurements. Round and long terminals were defined by W/H ratios (round = W/H ratio >0.2, long = W/H ratio <0.2). Data pooled from seven 3pw animals.

|  | Round |  |  |  |  | Long |  |  |  |  |
| --- | --- | --- | --- | --- | --- | --- | --- | --- | --- | --- |
| | n | Height<br>($\mu\text{m}$) | SE | Width<br>($\mu\text{m}$) | SE | n | Height<br>($\mu\text{m}$) | SE | Width<br>($\mu\text{m}$) | SE |
| Cervical | 31 | **145** | 28 | **49** | 11 | 45 | **259** | 56 | **28** | 5 |
| Thoracic | 3 | **155** | 6 | **37** | 6 | 81 | **334** | 61 | **24** | 4 |
| Lumbar | 99 | **157** | 20 | **48** | 5 | 83 | **274** | 32 | **28** | 3 |
| Sacral | 19 | **162** | 22 | **51** | 7 | 7 | **197** | 19 | **35** | 3 |

In the cervical and lumbar enlargements (C3-C6, L3-L6), the medial DH contains a curved zone of round arbors that is encircled by laterally located long arbors (Figure 4C). Somatotopic mapping of the cat and rat DH indicates that this medial curved zone in the lumbar enlargement matches the representation of the plantar paw and digits, with the dorsal paw and proximal limb representations lying more laterally^30, 31, 33-38^. We compared the number of labeled peripheral arbors in the toe and plantar paw skin with the number of round DH terminals in the corresponding half of the lumbar enlargement. This showed a very close correlation (Figure 4I), supporting that the round central arbors of the lumbar enlargement correspond to plantar paw and digit MrgprD^+^ nociceptors, while nociceptors from other regions of the hindlimb (including dorsal hindpaw and proximal hindlimb) grow long central arbors located more laterally.

Since <1% of MrgprD^+^ neurons are traced in these mice, it remains possible that this round-vs.-long arbor regional distinction may be an artifact of sparse labeling. For example, if both types were found throughout the DH but were differentially enriched between regions, sparse labeling might only trace the most prevalent type in each location. Using increasing dosages of tamoxifen, we found that these arbor types occupy mutually exclusive zones of the DH (Figure 5A-F). The maintained segregation of long and round arbors in the DH despite the increased number of AP^+^ DRG neurons indicates that these arbor morphologies represent a true somatotopic distinction among MrgprD^+^ non-peptidergic nociceptors.

**Figure 5.**
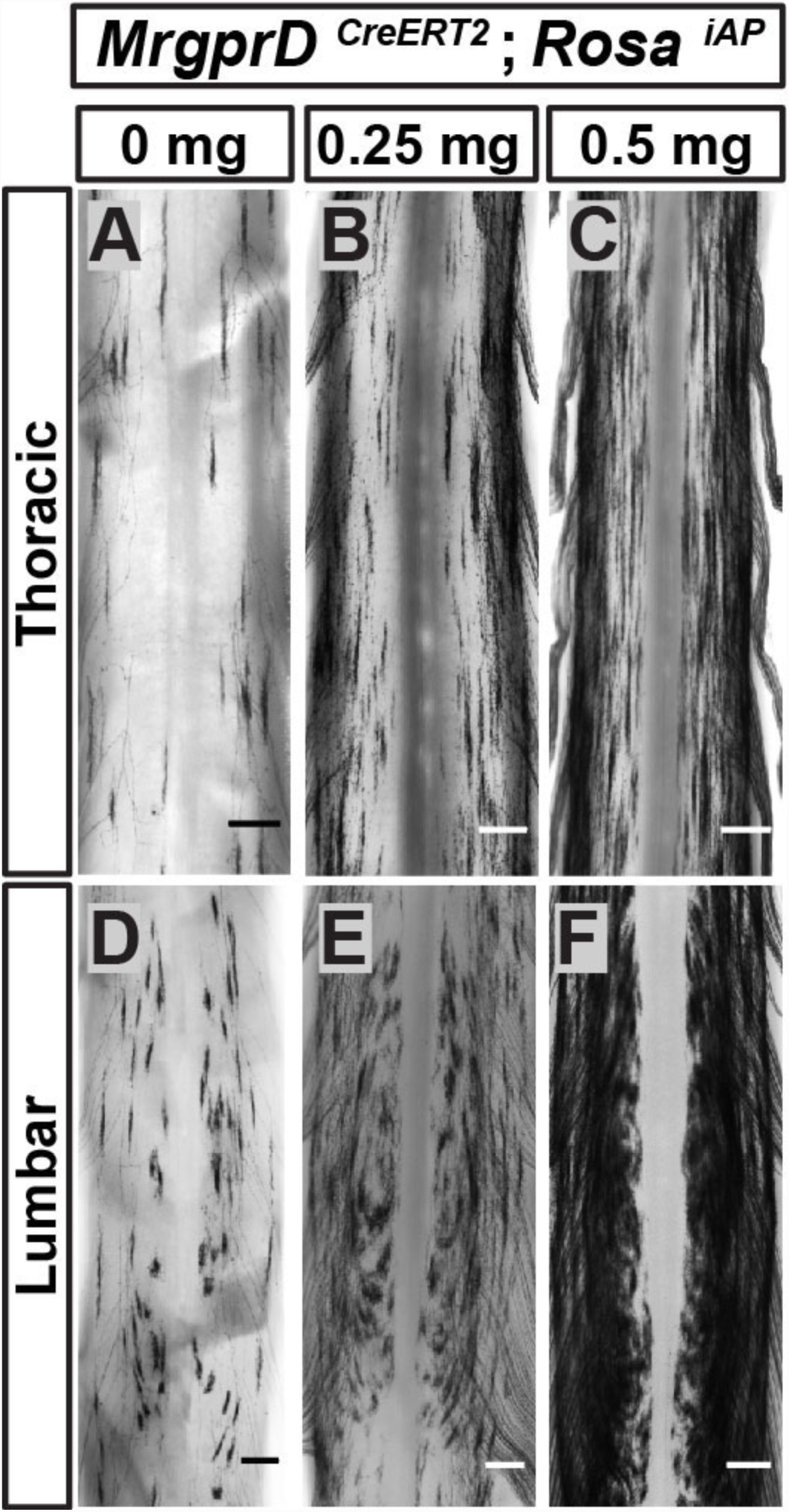
Non-peptidergic nociceptor labeling with increasing densities reveals somatotopic organization of MrgprD^+^ central arbors. (A-F) AP staining of *MrgprD^CreERT2^*; *Rosa^iAP^* thoracic (A-C) and lumbar (D-F) spinal cords that received prenatal 0mg (A&D), 0.25mg (B&E) or 0.5mg (C&F) prenatal tamoxifen. Even with increased labeling densities, round and long arbors occupy exclusive zones of the DH. *n =* 3 animals per treatment. Scale bars = 250μm.

### Neighboring non-peptidergic nociceptors highly overlap in the skin and spinal cord

The axonal arbors of some somatosensory neurons have a non-overlapping arrangement between neighbors (“tiling”) in the body wall of the fly and zebrafish^39, 40^. To determine if mammalian non-peptidergic nociceptive arbors tile in the skin, we generated double knock-in *MrgprD^CreERT2/EGFPf^*; *Rosa^tdTomato^* mice. After low-dose tamoxifen treatment, sparsely labeled MrgprD^+^ neurons in these mice express tdT while the entire population expresses EGFPf. The arbor fields of individual double tdT^+^/EGFP^+^ neurons are always co-innervated by EGFP^-^only^+^ fibers in both hairy and glabrous skin (Figure S5A-D), indicating that peripheral arbors of neighboring non-peptidergic nociceptors do not tile but instead overlap extensively. In hairy skin, sparse labeled tdT^+^ neurites co-innervated hair follicles with EGFP-only^+^ fibers, indicating that multiple MrgprD^+^ neurons can innervate single hair follicles. Similarly, both round and long DH arbors of non-peptidergic nociceptors highly overlap with their neighbors in the DH (Figure S5E&F).

### Heightened signal transmission in the paw DH circuitry of MrgprD+ neurons

Next, given the striking somatotopic differences in the central arbor organization of MrgprD^+^ nociceptors, we asked whether we could find regional (plantar paw vs. trunk) differences in the transmission of sensory information at the level of the dorsal horn. We generated *MrgprD^CreERT2^*; *Rosa^ChR2-EYFP^* mice, which were treated with postnatal tamoxifen (>P10, Figure 1), to compare the synaptic transmission of these neurons between somatotopic regions in spinal cord slice recordings. Comparable expression of ChR2-EYFP, as measured by the native fluorescence intensity, was found for MrgprD^+^ neurons in thoracic (T9-T12) and hind limb-level (L3-L5) DRGs (Figure S6). We first used *MrgprD^CreERT2^*; *Rosa^ChR2-EYFP/ChR2-EYFP^* mice (carrying two copies of the *Rosa^ChR2-EYFP^* allele) to perform *in vitro* whole-cell patch-clamp recordings of layer II interneurons located in the territory innervated by ChR2-EYFP^+^ fibers in transverse spinal cord sections (Fig. 6A). Light-evoked excitatory postsynaptic currents (EPSC_L_s) in these neurons could be differentiated into monosynaptic or polysynaptic responses based on latency, jitter, and response failure rate during 0.2 Hz blue light stimulation (Figure 6B&C)^41^. Almost all recorded neurons in both medial and lateral lumbar DH showed EPSC_L_s (Figure 6D, Table 4), with most cells (14/17 = 82.4% in medial lumbar, 14/16 = 86.5% in lateral lumbar) showing monosynaptic EPSC_L_s in both locations. This indicates that the majority of DH neurons in this innervation territory receive direct MrgprD^+^ input, and that the incidence receiving direct input is equivalent for medial lumbar and lateral lumbar DH.

**Table 4.**
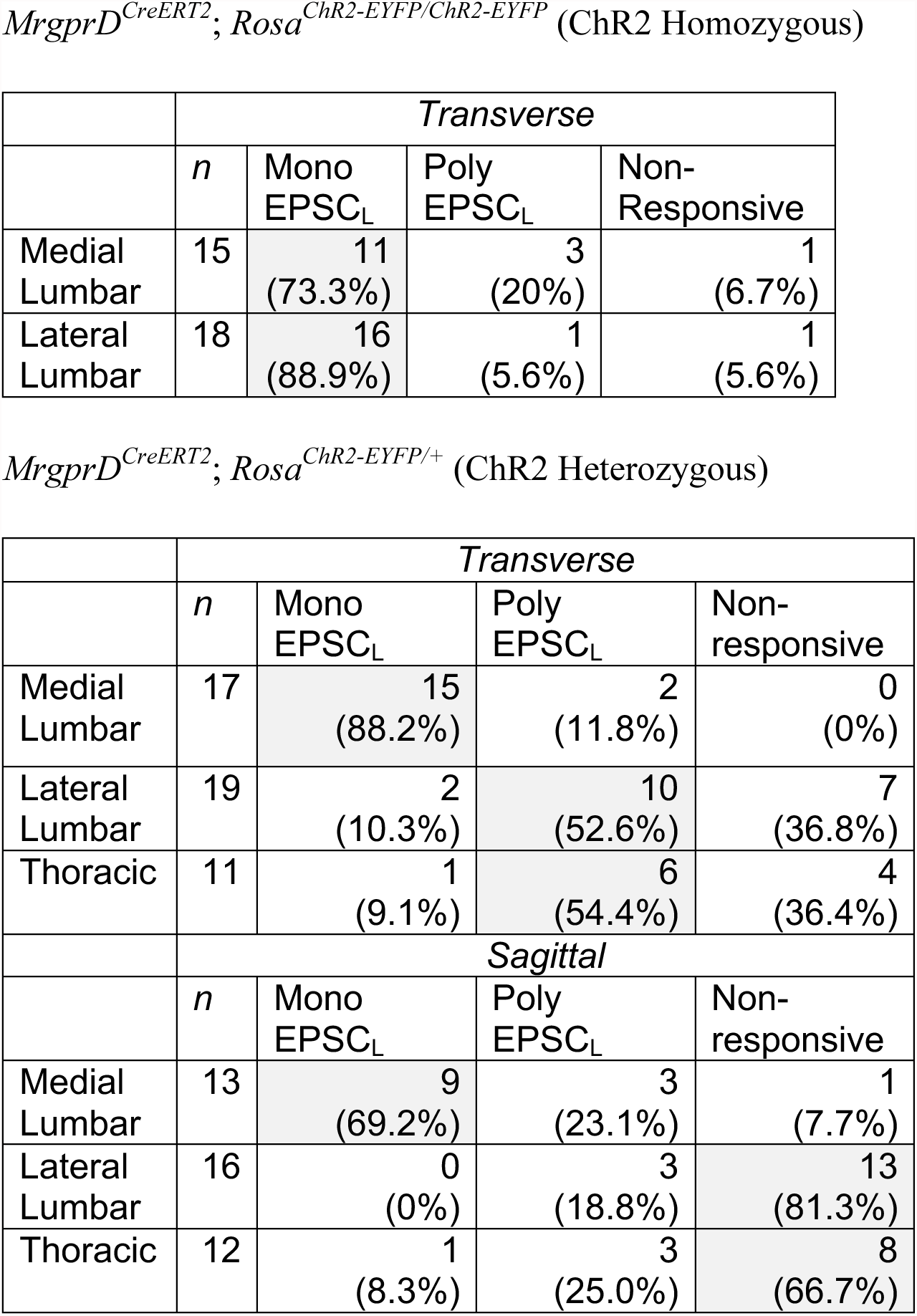
Summary of incidences of light-induced excitatory postsynaptic current (EPSC_L_) responses recorded from layer II neurons in *MrgprD^CreERT2^*; *Rosa^ChR2-EYFP^* homozygous and heterozygous mice. Patch clamp recordings were taken from either transverse or sagittal dSC slices, as indicated. Responses were classified as mono- or polysynaptic (see text). Shaded boxes show the response of the majority (>50%) of recorded cells.

Given the very high level of ChR2 expression in these mice, any potential difference between the medial and lateral lumbar spinal cord may be masked by a “ceiling” effect. We therefore halved the genetic dosage of *Rosa^ChR2-EYFP^* by taking slices from *MrgprD^CreERT2;^ Rosa^ChR2-EYFP/+^* mice. Interestingly, we saw a dramatic difference when comparing the medial and lateral lumbar DH of double heterozygous mice. Medial lumbar DH neurons showed a much higher incidence of light responses than lateral lumbar neurons, with a ~9-fold more (15/17 = 88.2% vs. 2/19 = 10.5%) in the incidence of monosynaptic EPSC_L_s over lateral lumbar neurons (Figure 6E, Table 4). Similar to the lateral lumbar DH, the incidence of monosynaptic EPSC_L_s of layer II neurons in the medial and lateral thoracic region are also very low (1/11 = 9.1%) (Figure 6E, Table 4). Moreover, even among responsive neurons, the pulse duration threshold required to elicit EPSC_L_s was much longer in lateral lumbar and thoracic DH neurons compared to medial lumbar neurons (Figure 6F). These results showed a lower threshold for light-trigged excitatory currents in medial lumbar compared to lateral lumbar and thoracic DH neurons, suggesting a heightened signal transmission in plantar paw circuits. We repeated these recordings in sagittal spinal cord slices and similarly found a higher EPSC_L_ incidence in medial lumbar compared to lateral lumbar or thoracic DH (Table 4). This confirms that these differences are not caused by the different orientation of nociceptors in round vs. long arbor regions. Taken together, our results show that, while most DH neurons in the MrgprD^+^ innervation territory receive direct MrgprD^+^ input, the overall signal transmission is heightened in plantar paw compared to trunk nociceptive circuits. This heightened transmission was seen specifically at the level of nociceptor-to-DH neuron connections, and it correlates closely with the region-specific organization of MrgprD^+^ central arbors.

**Figure 6.**
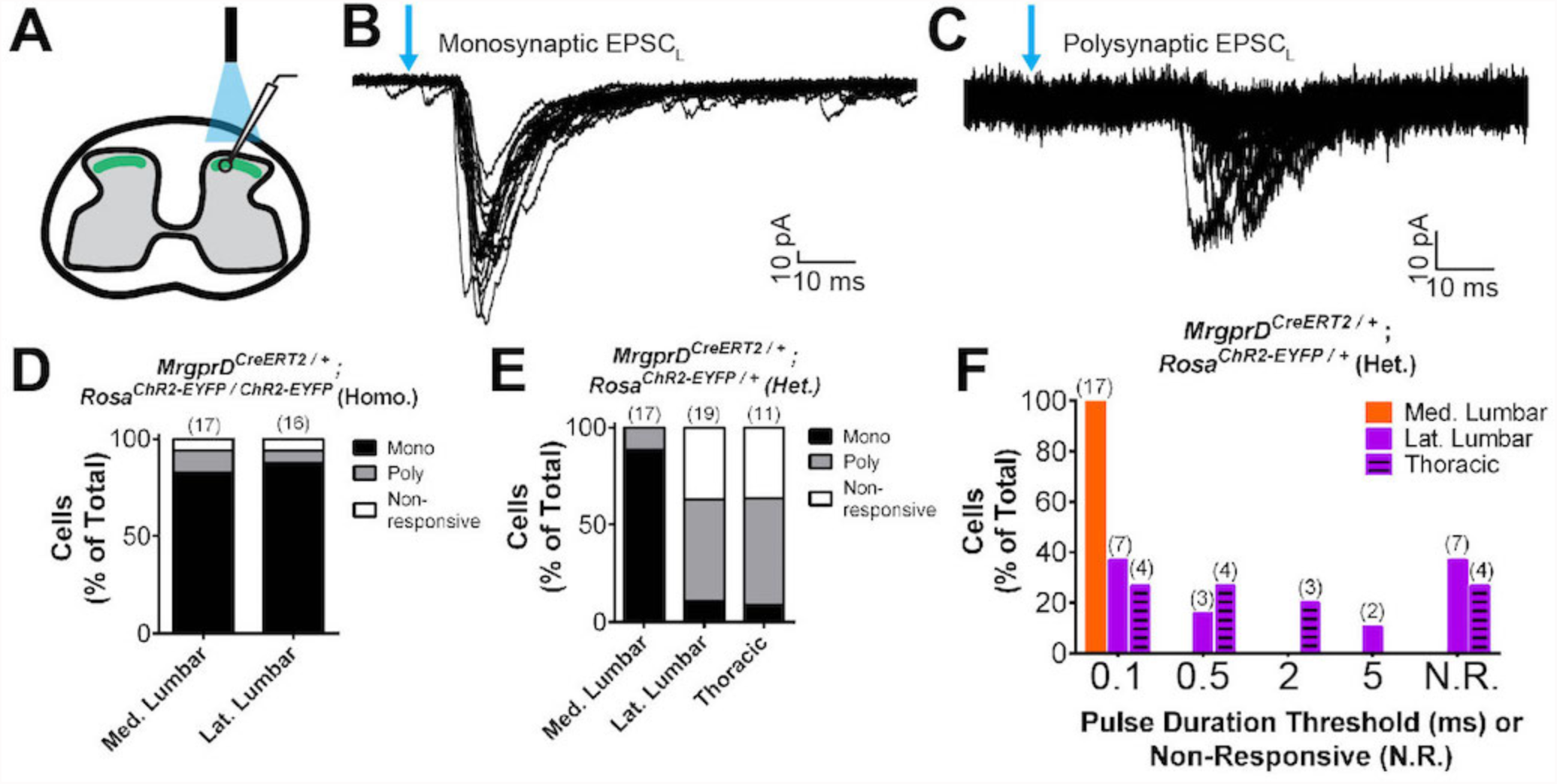
Plantar paw circuits show a heightened signal transmission in the dorsal horn. (A) Illustration of spinal cord slice recording from *MrgprD^CreERT2^*; *Rosa^ChR2-EYFP^* mice (P14-P21 tamoxifen) using optical stimulation. Neuron cell bodies located in the territory innervated by EYFP^+^ fibers were chosen for recording. (B&C) Monosynaptic (B) and polysynaptic (C) light-induced EPSC (EPSC_L_) traces recorded from layer II neurons during 0.2 Hz light stimulation (overlay of 20 traces). Light pulses indicated by blue arrows, scale bars shown in lower right. (D) In *MrgprD^CreERT2 / +^*; *Rosa^ChR2-EYFP/ChR2-EYFP^* homozygous slices, similar incidences of light-responsive neurons were found in medial and lateral lumbar regions. (E) In *MrgprD^CreERT2 / +^*; *Rosa^ChR2-EYFP/+^* heterozygous slices, a much higher incidence of light-responsive neurons was seen in medial lumbar compared to lateral lumbar or medial thoracic circuits. (F) Frequency distribution of threshold light pulse durations required for eliciting EPSC_L_s among cells in E. Among responsive cells, postsynaptic neurons in lateral lumbar and thoracic regions require longer pulse durations to be activated compared to those in medial lumbar region. Cell (*n*) numbers indicated in parentheses above bars in D-F.

### Plantar paw MrgprD^+^ nociceptors have a lower stimulation threshold to induce avoidance behaviors

Finally, since we saw a regional increase in signal transmission in plantar paw DH MrgprD^+^ circuits, we asked whether this has functional relevance for a freely behaving animal. To activate ChR2 in skin-innervating nociceptors of behaving mice, we stimulated *MrgprD^CreERT2^*; *Rosa^ChR2-EYFP/ChR2-EYFP^* (*MrgprD-ChR2*) and *Rosa^ChR2-EYFP/ChR2-EYFP^* (*Control*) mice (P10-17 tamoxifen treatment) at both the paw and upper-thigh leg skin (Figure 7B&E) with either 473nm blue light or 532nm green light as a negative control. High levels of ChR2-EYFP were expressed in peripheral neurites in both plantar paw and upper leg skin (Figure 7A&D). Consistent with the AP labeling (Figure 3), upper leg skin shows a much higher density of EYFP+ neurites than the paw glabrous skin (Figure S6A-C).

We first stimulated the paw skin of both groups of mice with green light (5mW, 10 Hz, sine wave) and observed no avoidance behavior such as paw withdrawal (Figure 7C, Videos 1&2). When we stimulated both groups of mice with 1mW of blue light (10 Hz, sine wave), control mice did not respond, while 12.5% of *MrgprD-ChR2* mice displayed light-induced paw withdrawal (Figure 7B&C). When stimulated with 5mW blue light (10 Hz, sine wave), 100% of *MrgprD-ChR2* mice displayed clear light-induced avoidance behavior responses consisting of paw withdrawal, high frequency paw shakes, occasional licking, extended holding of the paw in the air, and paw guarding. Control mice still did not respond (Figure 7B&C, Figure S7E, Videos 3&4).

**Figure 7.**
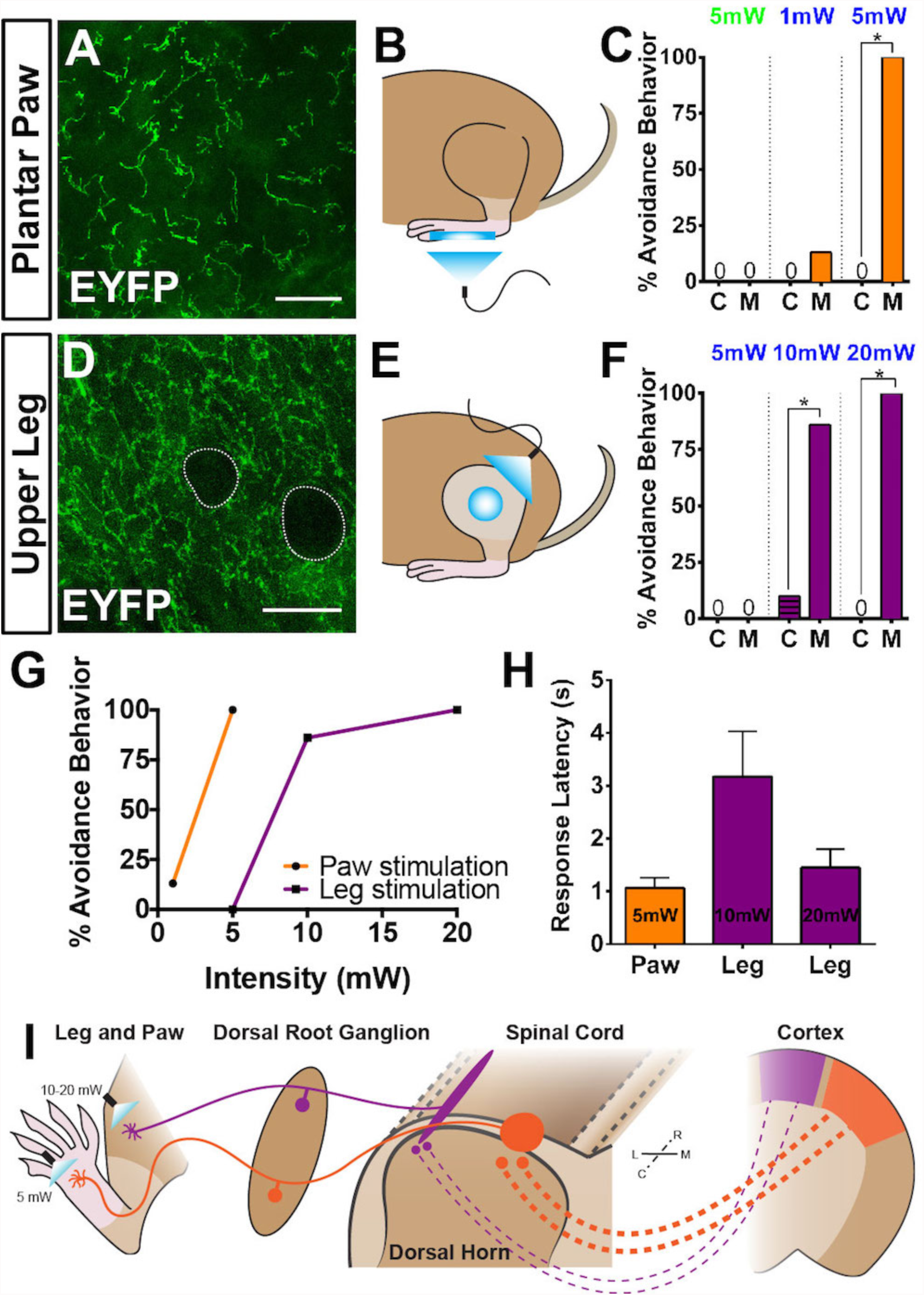
Peripheral optogenetic activation of MrgprD^+^ nociceptors reveals regional differences in optical threshold required to elicit withdrawal responses. (A-C) Optical stimulation in the paw. (A) Representative whole-mount immunostaining of plantar paw skin in *MrgprD^CreERT2 / +^*; *Rosa^ChR2-EYFP/ChR2-EYFP^* mice, *n*=3 mice. (B) Schematic of light placement on paw skin, see videos. (C) Histogram showing percentage of mice displaying aversive responses to 5mW green light and 1 or 5mW blue light to littermate control *(C)* and *MrgprD^CreERT2 / +^*; *Rosa^ChR2-EYFP/ChR2-EYFP^ (M). n* = 6-10 for each genotype with 1-2 trials per mouse. * = *p* <0.001 Chi-square test. (D-F) Optical stimulation in the leg. (D) Representative whole-mount immunostaining of upper leg skin in *MrgprD^CreERT2 / +^*; *Rosa^ChR2-EYFP/ChR2-EYFP^* mice, *n* = 3 mice. Dotted lines outline skin in *MrgprD^CreERT2 / +^*; *Rosa^ChR2-EYFP/ChR2-EYFP^* mice, *n* = 3 mice. Dotted lines outline hair follicles (E) Schematic of light placement on hair-shaven leg skin, see videos. (F) Histogram showing percentage of mice displaying aversive responses to 5, 10, or 20mW blue light at the leg (see above panel C for genotype description and statistical analyses). (G) A lower activation threshold is required for paw versus leg skin nociceptors. (H) Temporal delay time (seconds) from light onset to the first aversive behavior with 5, 10, or 20mW blue light in paw or leg of *MrgprD^CreERT2 / +^* ; *Rosa^ChR2-EYFP/ChR2-EYFP^* mice. Error bars represent SEM. (I) Model showing somatotopic organization of mammalian nociceptive circuitry. Distinct central arbor morphologies (“round versus long”) of MrgprD^+^ non-peptidergic nociceptors are observed in the medial versus lateral dorsal spinal cord, which correlates well with regional peripheral sensitivity and cortical representation. Scale bars = 50μm.

Strikingly, when we activated MrgprD^+^ neurites in the shaved upper-thigh skin, neither control nor *MrgprD-ChR2* mice responded to 5mW blue light (Figure 7E&F, S6E, Videos 5&6). Rather, to observe a fully penetrant avoidance behavior response in the upper-thigh of *MrgprD-ChR2* mice, the blue light power intensity had to be increased to 10 or 20 mW (2-4 times higher than the requirement in the paw) (Figure 7E-G, Videos 7&8). When taking the temporal delay of avoidance responses (Figure 7H) into consideration, 20mW intensity blue light is required at the leg to trigger responses comparable to 5mW intensity stimulation of the paw. In short, the light intensity threshold required to trigger an avoidance response is significantly lower in the paw compared to the upper-limb of *MrgprD-ChR2* mice.

Taken together, we have demonstrated a novel region-specific functional organization in the dorsal spinal cord for the mammalian pain system, which is consistent across anatomical, physiological, and behavioral levels.

## Discussion

In this study, we identified a novel somatotopic organization in the central arbors of mammalian MrgprD^+^ nociceptors, which is very well correlated with a regional increase in the sensitivity of paw nociceptive circuits to external stimuli. Our results suggest a model in which region-specific central arbor structure contributes to a sensory “fovea” for pain (Fig. 7I). In this model, the wider mediolateral spread of plantar paw nociceptor central arbors could facilitate “afferent magnification”^42^ in downstream CNS circuits and facilitate heightened pain sensitivity of the plantar paw. Remarkably, two features of mouse non-peptidergic nociceptors revealed by our study, the peripheral neurite density distribution and the nociceptive fovea location (the distal limb), are consistent with findings in humans^8-10^. Therefore, the organizational mechanisms we discovered in mice are likely to be conserved in human, which provides a possible explanation for the human “pain fovea”.

### *MrgprD^CreERT2^* allows for specific targeting of adult MrgprD^+^ nociceptors

Adult mice have two functionally distinct DRG neuronal populations expressing Mas-related gene product receptor (*Mrgpr*) genes. One expresses *MrgprA, MrgprB,* and *MrgprC* genes and the other only expresses *MrgprD*^23, 25, 43-47^. MrgprA/B/C^+^ neurons also transiently express *MrgprD* during early development ^25^. To specifically target MrgprD^+^ neurons, we generated a new inducible *MrgprD^CreERT2^* mouse line that allows for temporally controlled recombination. We demonstrated that postnatal (P10 or later) tamoxifen treatment of *MrgprD^CreERT2^* mice specifically targets MrgprD^+^ but not MrgprA/B/C^+^ neurons (Figure 1).

To our advantage, we also found that sparse recombination in untreated *MrgprD^CreERT2^*; *Rosa^iAP^* mice is very specific for MrgprD^+^ neurons (~94% AP^+^ neurons are MrgprD^+^, Figure 2I). We noticed that most of this random recombination likely occurs postnatally, as untreated *MrgprD^CreERT2^*; *Rosa^iAP^* tissue at P7 or younger shows no AP^+^ neurons (data not shown). This temporal delay of recombination, along with the fact that there are many more adult MrgprD^+^ than MrgprA/B/C^+^ neurons ^25, 43, 45, 46^, likely contributes to the specificity of sparse AP labeling in these mice.

In this paper, we used prenatal tamoxifen treatment of *MrgprD^CreERT2^* mice for two experiments: *MrgprD^CreERT2/EGFPf^*; *Rosa^tdTomato^* labeling to show nociceptor overlap (Figure S5) and increasing density labeling (Figure 5). In these experiments, both MrgprD^+^ and MrgprA/B/C^+^ neurons could be targeted. For the overlap experiment, neurites were chosen that had both red (*MrgprD^CreERT2^* ; *Rosa^tdTomato^*) and green (*MrgprD^EGFPf^*) fluorescence, indicating these were MrgprD^+^ neurites. For the increased density labeling experiment, even with high dosage (population-level) prenatal tamoxifen treatment of this line, MrgprA/B/C^+^ neurons make up <20% of the total cells labeled (Figure S2). Given the nature of this experiment, we believe that our interpretation is not confounded by this issue.

### Peripheral features of MrgprD^+^ nociceptors show no obvious mechanism for the sensory “fovea”

Though previous studies have traced various cutaneous somatosensory neurons, no systematic characterization of nociceptor morphology across the somatotopic map has been performed. In the periphery, sparse tracing of *Pou4f1* expressing somatosensory neurons (which includes almost all somatosensory DRG neuron classes^48^) in the back hairy revealed a “bushy ending” morphological type^29^. The authors suggested these terminals might correspond to C-fiber nociceptors or thermoceptors. In addition, MrgprB4^+^ and TH^+^ C fibers, which mainly mediate light touch but not pain, and innervate hairy skin only, have been genetically traced and analyzed^46, 49^. MrgprB4^+^ neurons innervate the skin in large patches of free terminals^46^, while TH^+^ neurons form lanceolate endings around hair follicles^49^. To our knowledge, single-cell tracing has not previously been performed for any C-fiber nociceptors innervating the glabrous skin.

MrgprD^+^ nociceptors innervate both hairy and glabrous skin but not deep tissues^24^, making this population ideal for analysis of somatotopic differences. Our *MrgprD^CreERT2^* tracing reveals that MrgprD^+^ nociceptors display a bushy-ending morphology in hairy skin and thickened endings in the epidermis in the glabrous skin (the plantar paw and finger tips) (Figure 3, Figure S3). Very rarely, we found “free ending” terminals that lack thickened epidermal ending (Figure S3). These match the morphology and size of MrgprB4^+^ light touch neurons^46^, possibly indicating very rare labeling of this subset. The fact that <2% of hairy skin terminals displayed this morphology further supports the specificity of *MrgprD^CreERT2^* sparse recombination.

Using genetic tracing, we found that (1) the density of MrgprD^+^ C fiber nociceptive neurites is slightly lower in paw (including digit tips) compared to trunk skin and (2) paw and trunk terminals are comparable in the overall area (Figure 3). This second result contrasts with an earlier study that performed single-fiber recordings of human mechanically responsive C-fiber units, which saw smaller receptive fields in the distal leg^11^. This discrepancy could be caused by differences between species, techniques (physiological vs. direct anatomic tracing), or the composition of neurons that were analyzed (the previous study presumably recorded from multiple molecular classes). In short, in our analysis of MrgprD^+^ nociceptors, we did not find any obvious peripheral mechanism that might readily explain the heightened pain acuity of the distal limbs seen in human subjects or the increased sensitivity of the mouse plantar paw to *in vivo* optogenetic skin stimulation (Figure 7).

### MrgprD^+^ nociceptors display region specific organization of central circuits

Classic studies have characterized the single-cell central arbors of C-fiber afferents using Golgi staining or backfill techniques and described them as longitudinally-oriented “thin sheets” that are short in the mediolateral axis and extended in the rostrocadual axis^18, 19, 50^. This description corresponds well to the “long arbors” we found in DH zones representing proximal limb and trunk regions (Figure 4). The central terminals of MrgprB4^+^ and TH^+^ C-fiber light touch neurons also match this long terminal morphology, consistent with the fact that these classes only innervate hairy skin^46, 49^. Nevertheless, a systematic comparison of nociceptive central terminals across the entire somatotopic map has not been conducted.

Here, we found regionally distinctive central arbor organization among MrgprD^+^ nociceptors; those innervating the distal limbs, tail, anogenital skin, and the head/face display round central terminals, while those innervating the trunk hairy skin display long and thin central arbors (Figure 4). Given that neurons in the medulla and sacral spinal cord display round arbors (Figure 4 and S4C), this morphological difference does not correlate with hairy vs. glabrous skin regions but instead seems to correlate with regions located at the extremities. In addition, we found that regions with round arbors showed an increased signal transmission onto downstream neurons, and a heighted sensitivity to external stimulation (Figures 6, 7, and S7). These results suggest that region specific central arbors of MrgprD+ nociceptors are an important component for the region-specific spinal cord circuits. Taken together, we have uncovered a novel form of region-specific functional organization for mammalian nociceptors.

Some previous studies suggest that the third-order DH collaterals of A**β** mechanoreceptors are also wider (in the mediolateral axis) in the medial lumbar enlargement compared to other DH regions^34, 51, 52^. However, the primary signal transmission for discriminative information occurs in the dorsal column nuclei but not the dorsal spinal cord. Thus, it is currently less clear whether this central arbor morphological difference contributes to differential touch sensitivity.

### Region-specific DH circuit organization leads to an increased sensitivity of paw pain circuits to external input

Finally, we determined the stimulus threshold (laser power) required to trigger avoidance behaviors of freely behaving *MrgprD-ChR2* mice upon paw or upper leg skin light stimulation. While 100% of tested animals showed withdrawal responses when 5 mW blue light was applied to the plantar paw, a 4-fold increase in light intensity (20 mW) was required to trigger comparable responses (in terms of rate and latency) in the upper leg (Figure 7, Figure S6B). Thus, our experiments show a clear heightened sensitivity of plantar-paw-innervating MrgprD^+^ nociceptors to stimulation. Interestingly, our use of peripheral optogenetic stimulation likely bypasses the effects of some peripheral parameters, such as the physical properties of the skin or the expression of molecular receptors, on the sensation of natural stimuli. Thus, this approach allowed us to more directly compare and probe the effects of circuit organization mechanisms on region-specific sensitivity.

Collectively, our results suggest that region-specific MrgprD^+^ central arbor organization could magnify the representation of the plantar paw within pain circuits to contribute to a nociceptive “fovea” ^52^ (Figure 7I). This central organization mechanism could allow for the sensitive detection of harmful stimuli in these areas, despite a lower neurite density in the periphery. This finding is relevant for pain research using mouse models, which has historically relied heavily on pain assays in plantar paw/medial lumbar circuits^53^. Given our findings, it would be interesting to examine whether region-specific differences exist in the molecular and physiological pathways of acute and/or chronic pain models. Such work could be informative for the translation of preclinical models to clinical treatment.

## Online Methods

### Mouse strains

Mice were raised in a barrier facility in Hill Pavilion, University of Pennsylvania. All procedures were conducted according to animal protocols approved by Institutional Animal Care and Use Committee (IACUC) of the University of Pennsylvania and National Institutes of Health guidelines. *Rosa^tdTomato^*, *Rosa^iAP^*, *Rosa^ChR2-EYFP^*, and *MrgprD^EGFPf^* lines have been described previously^24, 54-56^.

### Generation of *MrgprD^CreERT2^* mice

Targeting construct arms were subcloned from a C57BL/6J BAC clone (RP24-316N16) using BAC recombineering, and the CreERT2 coding sequence followed by a FRT-flanked neomycin-resistance selection cassette was engineered in-frame following the *MrgprD* starting codon by the same approach (Figure S1A). The targeting construct was electroporated into a C57/129 hybrid (V6.5) mouse embryonic stem cell line by the Penn Gene Targeting Core. ES clones were screened by PCR using primers flanking the 3’ insertion site (Figure S1C). Positive clones were further screened using Southern blot with both internal and external probes (Figure S1B). The Penn Transgenic Core assisted in *MrgprD^CreERT2^* ES clone blastocyst injection and in the generation of chimeric mice, which were mated to a *Rosa^Flippase^* line (Jax 007844) to excise the Neo cassette. Neo cassette-negative progeny (verified via PCR of genomic DNA) were mated to C57 or CD1 mice to establish the line.

### Genetic labeling of MrgprD^+^ nociceptors

To label MrgprD^+^ nociceptors, *MrgprD^CreERT2^* mice carrying the relevant reporter allele were treated with tamoxifen (Sigma, T5648) pre- or postnatally. For prenatal treatment, pregnant females were given tamoxifen along with estradiol (Sigma, E8875, at a 1:1000 mass estradiol: mass tamoxifen ratio) and progesterone (Sigma, P3972, at a 1:2 mass progesterone: mass tamoxifen ratio) in sunflower seed oil via oral gavage at E16.5-E17.5, when *MrgprD* is highly expressed in mouse non-peptidergic nociceptors^57^. For postnatal treatment, 0.5mg tamoxifen extracted in sunflower seed oil was given via i.p. injection once per day from P10-P17 (or P14-P21 for spinal cord slice recording experiments, Figure 6). At least one week was given to drive recombination and reporter gene expression.

### Tissue preparation and histology

Procedures were conducted as previously described^58^. Briefly, mice used for immunostaining or AP staining were euthanized with CO_2_ and transcardially perfused with 4% PFA/PBS, and dissected tissue (either skin or spinal cord and DRGs) was post-fixed for 2 hr in 4% PFA/PBS at 4° C. Tissue used for immunostaining was cryo-protected in 30% sucrose/PBS (4% overnight) before freezing. Mice used for *in situ* hybridization were euthanized and unfixed dissected tissue was frozen. Frozen glabrous skin and DRG/spinal cord sections (20-30 μm) were cut on a Leica CM1950 cryostat. Immunostaining of sectioned DRG, spinal cord, and glabrous skin tissue, whole mount skin immunostaining, and double fluorescence *in situ* hybridization was performed as described previously^58, 59^. The following antibodies and dyes were used: chicken anti-GFP (Aves, GFP-1020), rabbit anti-GFP (Invitrogen, A-11122), rabbit anti-NF200 (Sigma, N4142), rabbit anti-CGRP (Immunostar, 24112), conjugated IB4-Alex488 (Molecular Probes, I21411), chicken anti-peripherin (Aves, PER), chicken anti-PAP (Aves, PAP), mouse anti-PKCγ (Novex, 13-3800), rabbit anti-RET (IBL, 18121), rabbit anti-RFP (Chemicon, AB3216). *MrgprD*, *MrgprA3*, and *MrgprB4 in situ* probes were previously described^60^.

Tissue (skin or spinal cord with attached DRGs) for whole mount AP colorimetric staining with BCIP/NBT substrate (Roche, 1138221001 and 11383213001) and for fluorescent staining with HNPP/FastRed substrate (Roche, 11758888001) was treated as previously described^59^. Following AP colorimetric labeling, tissue was either cleared in BABB for imaging or sectioned using a VT1200S vibratome (Leica Microsystems, Nussloch, Germany) (200 μm), followed by BABB clearing for imaging. Fluorescent labeled DRGs co-stained using antibodies were cleared in glycerol and for imaging.

### Electrophysiology

Spinal cord slices recordings were conducted as previously described ^61^. Basically, 4-6pw *MrgprD^CreERT2^*; *Rosa^ChR2-EYFP/ChR2-EYFP^* or *MrgprD^CreERT2^* ; *Rosa^ChR2-EYFP/+^* mice were anesthetized with a ketamine/xylazine/acepromazine cocktail. Laminectomy was performed, and the spinal cord lumbar segments were removed and placed in ice-cold incubation solution consisting of (in mM) 95 NaCl, 1.8 KCl, 1.2 KH_2_PO_4_, 0.5 CaCl_2_, 7 MgSO_4_, 26 NaHCO_3_, 15 glucose, and 50 sucrose, oxygenated with 95% O_2_ and 5% CO_2_, at a pH of 7.35–7.45 and an osmolality of 310–320 mosM. Sagittal or transverse spinal cord slices (300-500 μm thick) were prepared using a VT1200S vibratome (Leica Microsystems, Nussloch, Germany) and incubated in 34°C incubation solution for 30 min.

The slice was transferred to the recording chamber and continuously perfused with recording solution at a rate of 3–4 ml/min. The recording solution consisted of (in mM) 127 NaCl, 1.8 KCl, 1.2 KH_2_PO_4_, 2.4 CaCl_2_, 1.3 MgSO_4_, 26 NaHCO_3_, and 15 glucose, oxygenated with 95% O_2_ and 5% CO_2_, at a pH of 7.35–7.45 and an osmolality of 300–310 mosM. Recordings were performed at RT. Spinal cord slices were visualized with an Olympus BX 61WI microscope (Olympus Optical, Tokyo, Japan), and the substantia gelatinosa (lamina II), which is a translucent band across the dorsal horn, was used as a landmark. Fluorescently labeled ChR2-EYFP terminals in the DH were identified by epifluorescence, and neurons in this innervation territory were recorded in the whole cell patch-clamp configuration. Glass pipettes (3–5 MΩ) were filled with internal solution consisting of (in mM) 120 K-gluconate, 10 KCl, 2 MgATP, 0.5 NaGTP, 20 HEPES, 0.5 EGTA, and 10 phosphocreatine di(tris) salt at a pH of 7.29 and an osmolality of 300 mosM. All data were acquired using an EPC-9 patch-clamp amplifier and Pulse software (HEKA, Freiburg, Germany). Liquid junction potentials were not corrected. The series resistance was between 10 and 25 MΩ.

Light induced EPSCs (EPSC_L_s) were elicited at a frequency of 2/min by 473nm laser illumination (10 mW, 0.1-5ms, Blue Sky Research, Milpitas, USA). Blue light was delivered through a 40X water-immersion microscope objective. Mono- or polysynaptic EPSC_L_s were differentiated by 0.2 Hz light stimulation. We classified a connection as monosynaptic if the EPSC jitter (average standard deviation of the light-induced EPSCs latency from stimulation) < 1.6 ms^62, 63^.

### Optogenetic stimulation of MrgprD^+^ nociceptors in paw and leg skin

To induce light-evoked behavior in freely moving animals, we used P10-P17 tamoxifen treated *MrgprD^CreERT2^; Rosa^ChR2-EYFP/ChR2-EYFP^* mice and control littermates (*Rosa^ChR2-EYFP/ChR2-EYFP^*) who were also tamoxifen treated, but lacked the Cre-driver. All tested animals were between 2-6 months old. An additional control was to shine 532nm green laser light (Shanghai Laser and Optics Century, GL532T8-1000FC/ADR-800A) to the paw skin of both experimental groups. To induce light-evoked aversive behavior in MrgprD^+^ neurites in the paw skin, 1 or 5mW of 473nm blue light laser (Shanghai Laser and Optics Century, BL473T8-150FC/ADR-800A) was shined directly to the paw skin through a mesh bottom floor, with a cutoff time of 10 seconds. We tested different light waveforms and found that 10Hz sine waveform pulsing gave the best behavior responses. Thus, we used this waveform for all our behavior tests.

To induce light-evoked aversive behavior in MrgprD^+^ neurites in the leg skin, first, all hair was removed from the leg with Nair hair remover under light 3% isoflourane anesthesia, and animals were given two days before being tested again. For leg stimulation, 1, 5, 10, or 20mW of 473nm blue light laser, with 10Hz sine waveform pulsing, was shined directly to the leg skin. The cutoff time for this behavior assay is 10 seconds. For habituation to either paw or leg skin light stimulation, on the first two days, animals were habituated to the testing paradigm by being placed in the plexiglass testing chamber (11.5×11.5×16 cm) for 30 minutes each day. On the third testing day, animals were placed in the plexiglass testing chamber for 15 minutes prior to light stimulation. Green and blue light testing were performed on different days, but two weeks separated the paw and leg skin light stimulation. For all stimulation, the laser light was delivered via an FC/PC Optogenetic Patch Cable with a 200micrometer core opening (ThorLabs) and there was approximately 1cm of space between the cable terminal and the targeted skin area. Light power intensity for each experiment was measured with a Digital Power Meter with a 9.5mm aperture (ThorLabs). For leg skin stimulation, power intensity was only slightly impeded by the thin wall (0.02cm) of the plexiglass holding chamber, as measured by the Digital Power Meter (Figure 7D).

To gain precise spatial and temporal resolution of behavior responses, we recorded behaving animals at 500 frames/second with a high-speed camera (FASTCAM UX100 800K-M-4GB - Monochrome 800K with 4GB memory) and attached lens (Nikon Zoom Wide Angle Telephoto 24-85mm f2.8). With a tripod with geared head for Photron UX100, the camera was placed at a ~45degree angle at ~1-2 feet away from the plexiglass holding chambers where mice performed behaviors. The camera was maximally activated with far-red shifted 10mW LED light that did not disturb animal behavior. All data were collected and annotated on a Dell laptop computer with Fastcam NI DAQ software that is designed to synchronize Photron slow motion cameras with the M series integrated BNC Data Acquisition (DAQ) units from National Instruments.

### Image acquisition and data analysis

Images were acquired either on a Leica DM5000B microscope (bright field with a Leica DFC 295 camera and fluorescent with a Leica 345 FX camera), on a Lecia SP5II confocal microscope (fluorescent), or on a Leica M205 C stereoscope with a Leica DFC 450 C camera (bright field). Image manipulation and figure generation were performed in ImageJ, Adobe Photoshop and Adobe Illustrator. Cell number counting, nociceptor arbor measurements, and fluorescence measurements were performed in ImageJ. For section histology experiments, *n* = 3-6 sections per animal from 3 animals. Central arbor height/width measurements were taken to be the relevant axes of fitted ellipses. Column graphs, pie charts and scatter plots were generated in GraphPad Prism5. Column graphs show mean ± SEM. Statistical significance was analyzed using Student’s *t*-tests, one-way ANOVA with Tukey’s multiple comparisons or Chi-square tests in GraphPad Prism5. For DRG fluorescence intensity measurements, the average fluorescence of outlined cells was normalized to background fluorescence (Figure S6).

## Acknowledgements

We thank Peter Dong for illustrating the model in Figure 7. We also thank Greg Bashaw for reading and providing suggestions for this manuscript. This work was supported by National Institutes of Health (NIH) grant (NS083702, NS094224) and the Klingenstein-Simons Fellowship Award in the Neurosciences to W.L. and by NIH grant (NS092297) to W.O. I.A.S. was supported by NIH grant (K12GM081259) and Burroughs Wellcome Fund grant PDEP.

## Author Contributions

W.O., I.A.S., L.C., M.M. and W.L. contributed to experimental design and interpretation. W.O. performed most of the histological experiments in this paper. L.C. performed the spinal cord slice recording experiments. I.A.S. and J.B. performed the optogenetic behavior assays. W.O., I.A.S., L.C., M.M. and W.L. wrote and revised this manuscript.

## Supplemental Figures and Videos

**Supplementary Figure S1.**
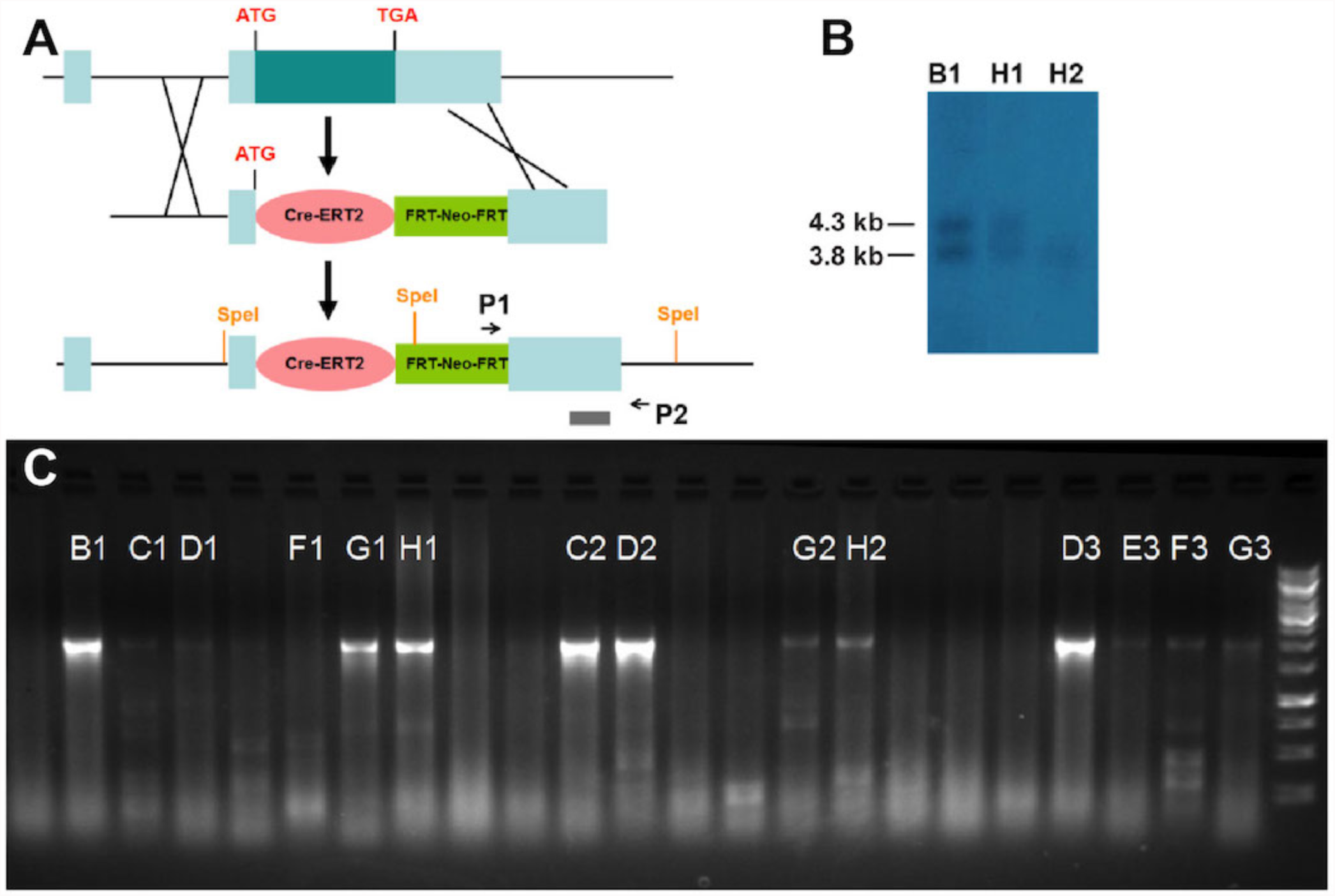
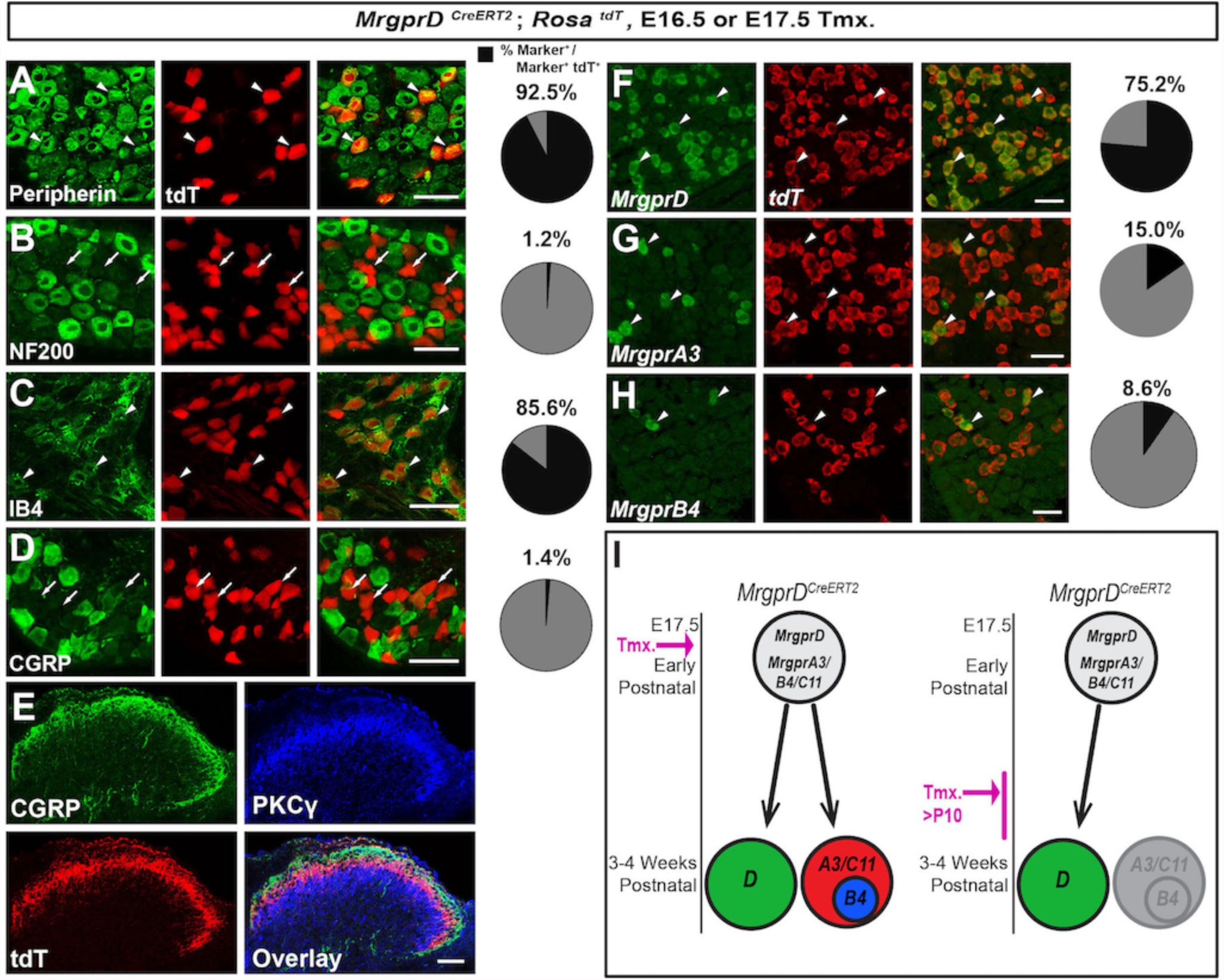
Generation of *MrgprD^CreERT2^* knock-in mouse line. (A) Illustration showing the knock-in targeting strategy. Grey bar, 3’ UTR Southern blot probe site. P1, P2, primers for PCR screening. (B) Southern blot of SpeI-digested ES genomic DNA, using a probe against *MrgprD* 3’ UTR (grey bar in A). 4.3 kb, *MrgprD^CreERT2^* knock-in allele. 3.8 kb, *MrgprD* wild-type allele. (C) PCR screen of electroporated ES clones, using P1, P2 primers. The positive PCR product is about 2 kb.

**Supplementary Figure S2.**
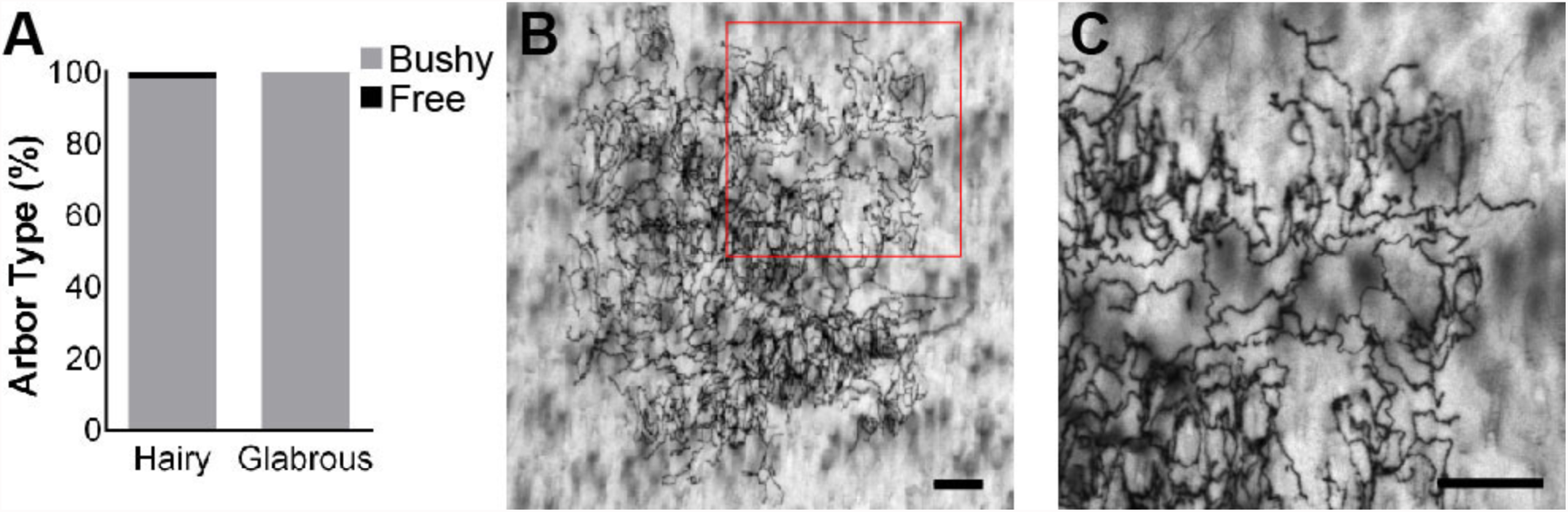
Prenatal tamoxifen treatment labels MrgprD^+^ along with MrgprA3/B4^+^ non-peptidergic DRG neurons. (A-D) Representative DRG sections of 3pw *MrgprD^CreERT2^*; *Rosa^tdTomato^* mice (2.5-5 mg tamoxifen at E16.5) immunostained with the indicated markers. tdT^+^ neurons are positive for non-peptidergic neuron markers peripherin (92.5 ± 4.0%) (A) and IB4 (85.6 ± 1.2%) (C), but do not express NF200 (1.2 ± 0.01%) (B) or CGRP (1.4 ± 0.00%) (D). (E) Representative immunostained DH section showing tdT^+^ fibers innervating layer II, ventral to CGRP^+^ fibers but dorsal to PKCγ interneurons. (F-H) Double fluorescent *in situ* shows expression of *MrgprD* (75.2 ± 0.8%) (F), *MrgprA3* (15.0 ± 0.2%) (G) and *MrgprB4* (8.6 ± 0.1%) (H) in *tdT* expressing neurons from prenatally treated *MrgprD^CreERT2^*; *Rosa^tdTomato^* mice (5 mg tamoxifen at E17.5). (I) Model showing *MrgprD^CreERT2^* specificity based on time of tamoxifen dosage. *MrgprD^CreERT2^* recombination is consistent with expression of *MrgprD* across development. Arrowheads indicate marker overlap with tdT, arrows indicate tdT^+^ cells that do not express indicated marker. Scale bars, 50μm (A-D, F-H), 100μm (E).

**Supplementary Figure S3.**
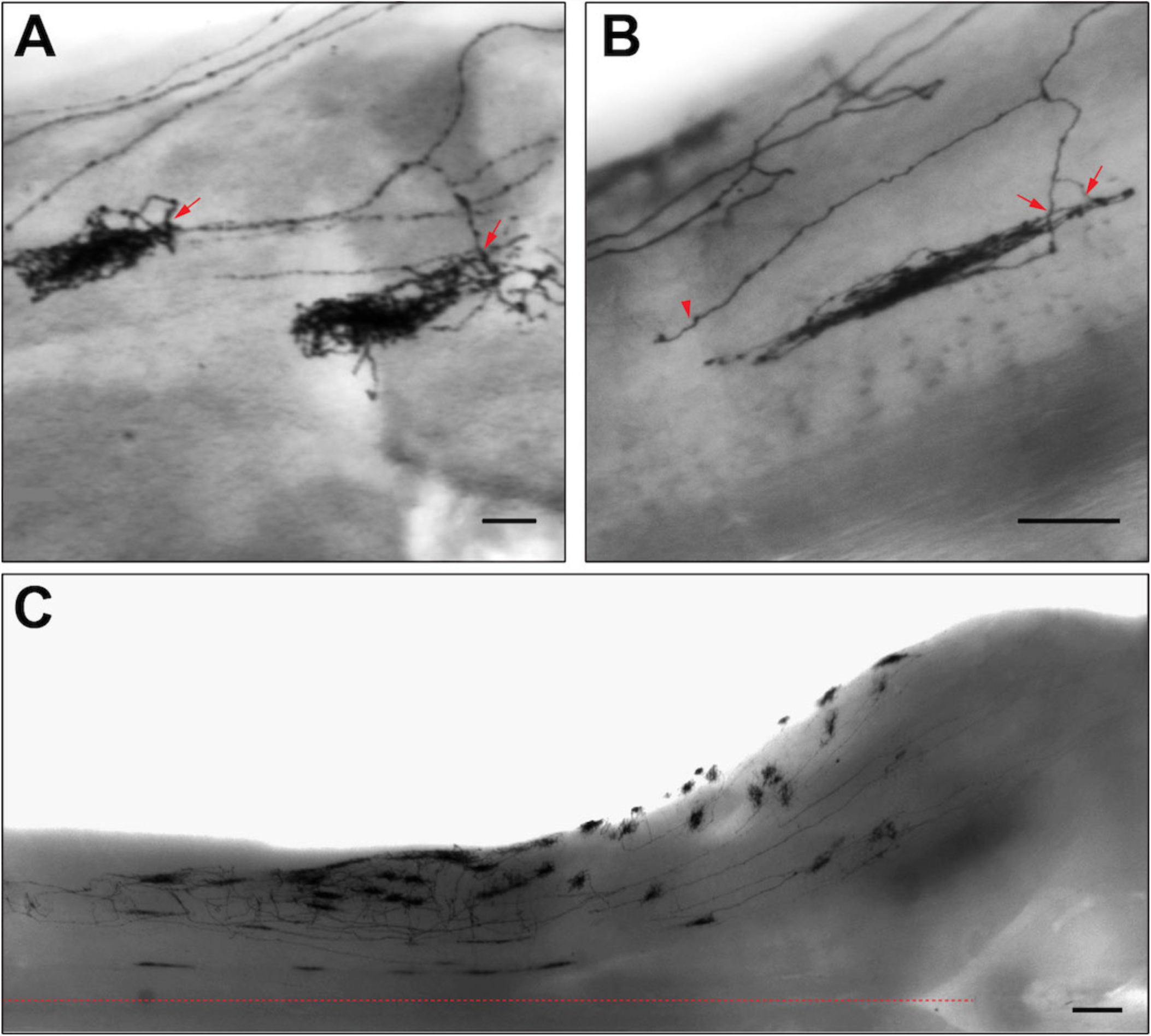
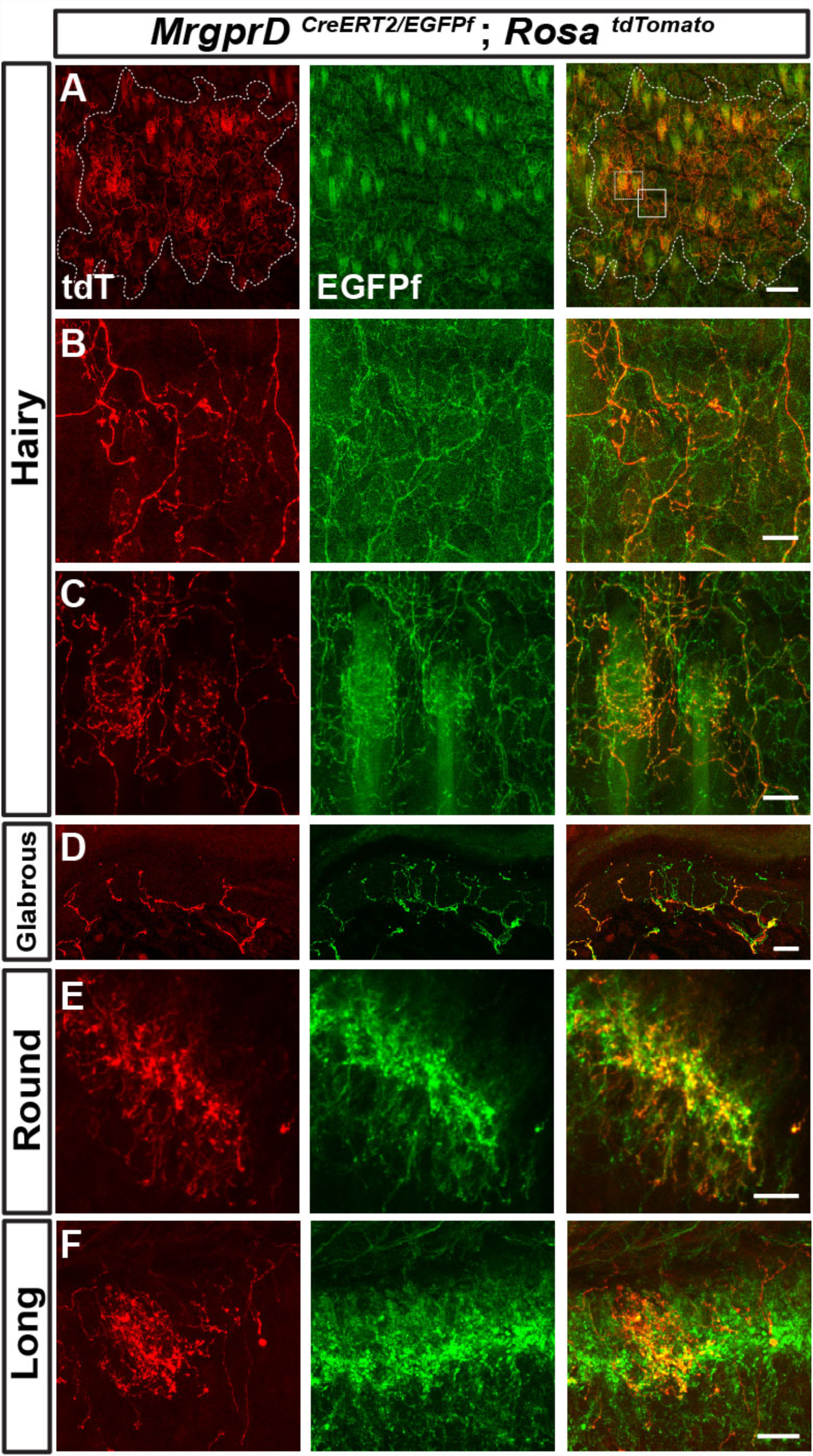
Rare free arbors are found in sparsely labeled non-peptidergic nociceptors of *MrgprD^CreERT2^* ; *Rosa^iAP^* mice. (A) Percentage of bushy or free ending type arbors by location. *n* = 132 hairy, 22 glabrous terminals from 4 animals. (B&C) Sparse labeled free terminal-type arbor in trunk hairy skin. C is a high magnification view of the region boxed in B. Scale bars = 100μm.

**Supplementary Figure S4**. Central arbors of sparsely labeled non-peptidergic neurons in *MrgprD^CreERT2^*; *Rosa^iAP^* mice. (A&B) Examples of bifurcating non-peptidergic nociceptor central projections from 1pw (0.05 mg tamoxifen at E16.5) (A) and 3pw (B) spinal cords. Bifurcated branches sometimes give rise to independent terminals (arrows in A), join other branches to give rise to a common terminal (arrows in B) or end without elaborating a terminal (arrowhead in B). (C) Round terminals in the upper cervical spinal cord and medulla, many of which descend from the TG non-peptidergic neurons. Scale bars, 50μm (A&B), 250μm (C).

**Supplemental Figure S5.**
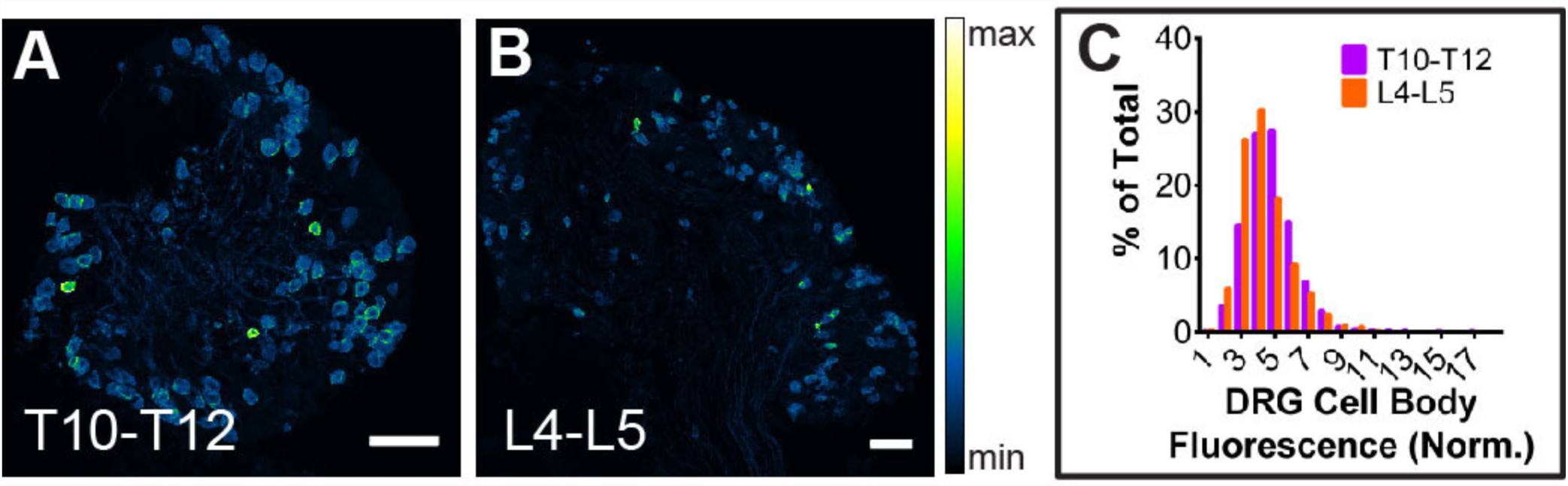
Neighboring non-peptidergic nociceptors overlap extensively in the skin and spinal cord. (A-C) Whole mount immunostaining of *MrgprD^CreERT2/EGFPf^*; *Rosa^tdTomato^* (0.5 mg tamoxifen at E16.5) hairy skin with anti-GFP and anti-RFP antibodies. The terminal field of one non-peptidergic nociceptor is labeled with tdT, as outlined in A. B&C show higher magnifications view of the regions boxed in A (solid line = B, dotted line = C). Innervation of hair follicles is shown in C. (D) Immunostaining of a section of glabrous skin. (E&F) Immunostaining of medial cervical (D) and thoracic (E) spinal cord sections, showing sparse labeled round terminal and long terminals. Scale bars = 100μm (A), 20μm (B-F).

**Supplementary Figure S6.**
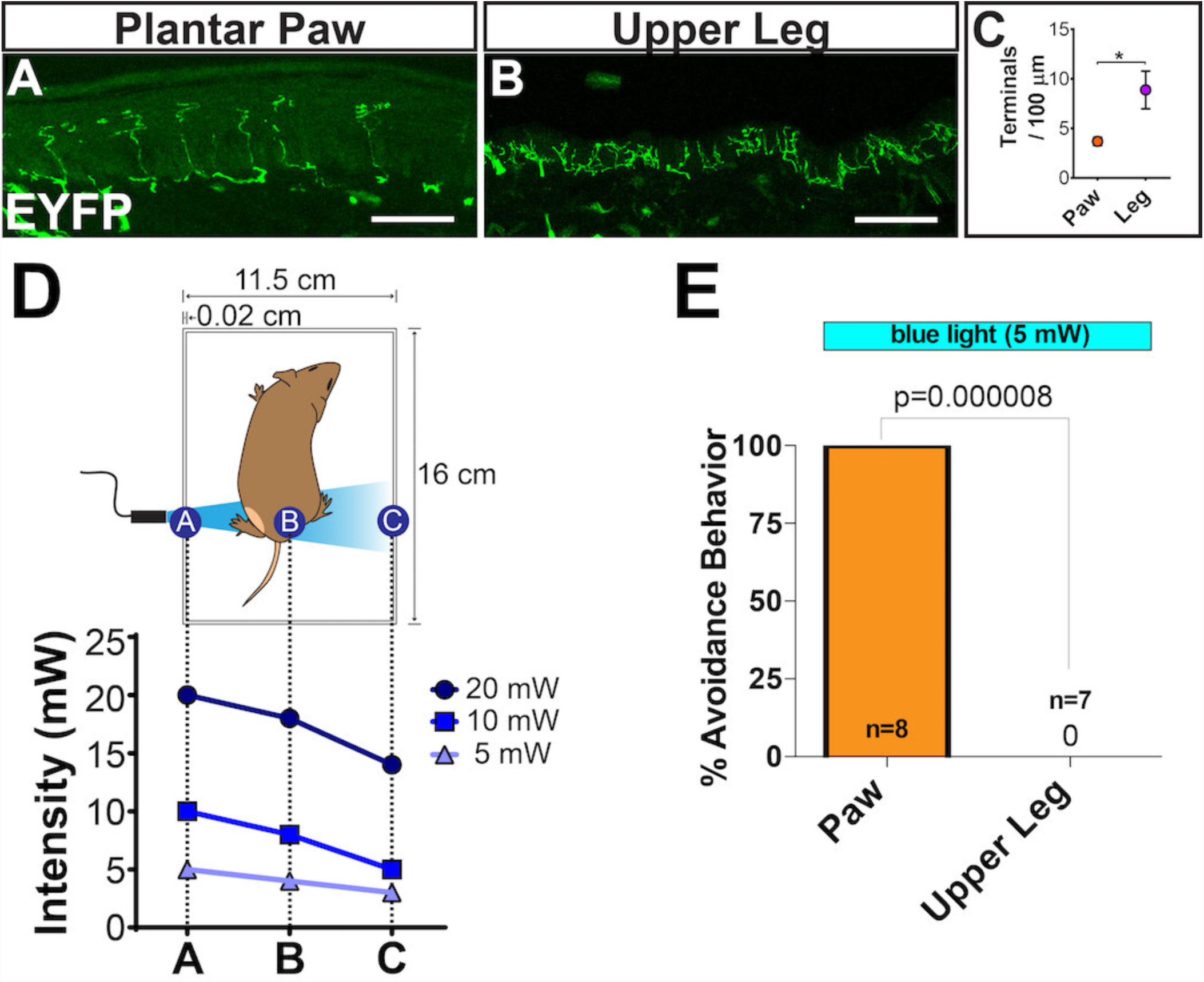
MrgprD^+^ neurons show no regional difference in *Rosa^ChR2-EYFP^* expression. (A&B). Native (no immunostaining) ChR2-EYFP fluorescence of *MrgprD^CreERT2^*; *Rosa^ChR2-EYFP^* (P10-P17 tamoxifen) DRG sections. (C) T10-T12 and L4-L4 DRG cell body fluorescence frequency distributions overlap, indicating no regional change in ChR2-EYFP expression level. Scale bars = 100μm.

**Supplementary Figure S7**. *In vivo* optogenetic peripheral stimulation. (A-C) *MrgprD^CreERT2^*; *Rosa^ChR2-EYFP^* sectioned upper leg hairy skin (B) shows a higher density of ChR2-EYFP^+^ neurites than plantar paw glabrous skin (A). (C) Quantification of neurites per 100μm. *n* = 3 animals. * = *p* <0.05, Chi-square test. (D) Measured light power at three locations (closest side, midpoint, farthest side) in the behavior chamber with 5 mW, 10 mW, and 20 mW blue laser intensity. (E) Response rate (% of mice) showing withdrawal responses to 5 mW blue light stimulation at paw or upper leg. *n* = 6-10 mice, 1-2 trials per mouse. Scale bars = 50 μm (A&B).

**Supplementary Video 1**. High speed video recording of green laser light (5 mW) plantar paw stimulation of Control (*ChR2*) mice. Green light does not induce an avoidance response.

**Supplementary Video 2**. High speed video recording of green laser light (5 mW) plantar paw stimulation of *MrgprD*; *ChR2* mice. Green light does not induce an avoidance response.

**Supplementary Video 3**. High speed video recording of blue laser light (5 mW) plantar paw stimulation of Control mice. Blue light does not induce an avoidance response in Control mice.

**Supplementary Video 4**. High speed video recording of blue laser light (5 mW) plantar paw stimulation of *MrgprD*; *ChR2* mice. 5 mW blue laser light triggers robust avoidance responses, including paw withdrawal and shaking, when applied to the paw of *MrgprD*; *ChR2* mice.

**Supplementary Video 5**. High speed video recording of blue laser light (5 mW) shaved upper leg stimulation of Control mice. No avoidance response is seen.

**Supplementary Video 6**. High speed video recording of blue laser light (5 mW) upper leg stimulation of *MrgprD*; *ChR2* mice. 5 mW blue laser light is insufficient to trigger avoidance responses when applied to the shaved upper leg of *MrgprD*; *ChR2* mice.

**Supplementary Video 7.** High speed video recording of blue laser light (20mW) upper leg stimulation of Control mice. No avoidance response is seen, indicating that 20 mW laser light stimulation is not by itself aversive.

**Supplementary Video 8.** High speed video recording of blue laser light (20mW) upper leg stimulation of *MrgprD*; *ChR2* mice. 20 mW laser light induces robust avoidance behavior, including limb withdrawal and licking, when applied to the shaved upper leg of *MrgprD*; *ChR2* mice.

